# Systems-informed genome mining for electroautotrophic microbial production

**DOI:** 10.1101/2020.12.07.414987

**Authors:** Anthony J. Abel, Jacob M. Hilzinger, Adam P. Arkin, Douglas S. Clark

## Abstract

Microbial electrosynthesis (MES) systems can store renewable energy and CO_2_ in many-carbon molecules inaccessible to abiotic electrochemistry. Here, we develop a multiphysics model to investigate the fundamental and practical limits of MES enabled by direct electron uptake and we identify organisms in which this biotechnological CO_2_-fixation strategy can be realized. Systematic model comparisons of microbial respiration and carbon fixation strategies revealed that, under aerobic conditions, the CO_2_ fixation rate is limited to <6 μmol/cm^2^/hr by O_2_ mass transport despite efficient electron utilization. In contrast, anaerobic nitrate respiration enables CO_2_ fixation rates >50 μmol/cm^2^/hr for microbes using the reductive tricarboxylic acid cycle. Phylogenetic analysis, validated by recapitulating experimental demonstrations of electroautotrophy, uncovered multiple probable electroautotrophic organisms and a significant number of genetically tractable strains that require heterologous expression of <5 proteins to gain electroautotrophic function. The model and analysis presented here will guide microbial engineering and reactor design for practical MES systems.

## Main

The capture and conversion of CO_2_ to fuels, commodity chemicals, and pharmaceutical precursors can help close the anthropogenic carbon cycle. Biological CO_2_ fixation using plants, algae and cyanobacteria occurs naturally at scale, but biotechnological application of photosynthetic carbon fixation is challenging for several reasons including low conversion rates, low efficiencies, and difficulties with downstream separations^1^. Physicochemical strategies to fix CO_2_ by generating syngas have also been considered, but extreme operating conditions and low product selectivity for complex hydrocarbons have hindered adaptation^2–4^. Recently, electromicrobial approaches, in which electrochemical reduction of CO_2_ provides C_1_ feedstocks to biochemical processes producing a wide array of chemicals, have been proposed^5–7^. This strategy is particularly promising because it benefits from extensive system modeling and abiotic catalyst discovery efforts^8–11^. However, poor catalyst stability, reliance on rare elements, and potential catalyst toxicity could inhibit the scalability of this method^12–14^. Microbial electrosynthesis (MES) systems may overcome these issues by obviating the need for an electrocatalyst by using so-called electroautotrophic microbes that accept electrons from a cathode via direct electron transfer (DET) mechanisms^15,16^. The inherent regenerative capacity of microbes also makes MES an attractive option for chemical production during space exploration missions because carry-along mass and materials resupply challenges are key constraints on long-term or deep-space expeditions.^17^

Despite the promise of direct MES, product spectrum and production rate bottlenecks have prevented technological realization^18^. MES systems developed to date primarily use acetogenic or methanogenic microbes that divert most of their fixed carbon into low-value acetate and methane^19^. Although genetic tools have recently become available for some of these organisms^20,21^, microbial energy conservation strategies severely restrict the achievable product spectrum and selectivity. Microbes supporting electroautotrophy via the Calvin cycle have recently been discovered and are likely to alleviate product spectrum issues^22,23^. However, high throughput platforms for discovery and engineering of novel microbial chassis are in their nascent stages and distinguishing between more and less promising candidates and identifying engineering targets that enable high production rates is challenging^24,25^.

Computational MES models can address this challenge by comparing microbial respiration and carbon fixation strategies. To that end, several models of MES systems have been developed, but these have assumed electron uptake interfaces directly with the intracellular NAD^+^/NADH pool, in contrast to known electron transfer mechanisms^26,27^. Recent energetic calculations to determine the limiting efficiency of MES systems have addressed this issue, but assumed that aerobic respiration is equally available for all carbon fixation pathways (CFPs)^28^. Moreover, considerations of physiological mechanisms of electron transfer, respiration, and carbon fixation in models that capture relevant physical phenomena remains an outstanding challenge.

Here, we incorporate a physiological, mechanistic understanding of extracellular electron uptake into a comprehensive multiphysics model of MES that describes mass transport, electrochemical and acid-base thermodynamics and kinetics, and gas-liquid mass transfer. In the proposed mechanism, based on the reversible electron conduit in *Shewanella oneidensis*^29^, electrons supplied by the cathode are split in a bifurcation scheme. A fraction is used to produce a proton motive force (PMF) via aerobic or anaerobic nitrate respiration, while the remainder is used along with the PMF to regenerate cellular energy carriers (ATP, NAD(P)H, reduced ferredoxin) consumed in the CFPs. Using this detailed picture of electron uptake, we compare the productivity and efficiency of MES systems with hypothetical microbes performing carbon fixation with each of four major CFPs and we identify physiological modules that enable the highest productivities. We further perform phylogenetic analysis of marker genes for these modules to uncover naturally occurring and/or readily engineerable microbial chassis that require the heterologous expression of only a few proteins. Thus, our analysis provides crucial insight into microbial catalyst discovery and engineering and reactor design strategies that can advance direct MES systems from basic science to technological practice.

## Results

### System overview

The MES model (see Computational Methods) considers a one-dimensional bioelectrochemical reactor for CO_2_ reduction (Fig. 1a). The reactor has well-mixed anolyte and catholyte regions that are replenished at a fixed dilution rate and to which CO_2_ is constantly supplied at a fixed partial pressure. These regions are separated by an anion exchange membrane (AEM) and fluid boundary layers, which also separate the well-mixed phases from the anode surface and the biocathode layer. The chemical species in each chamber are dissolved CO_2_, dissolved O_2_ (in the case of aerobic operation), bicarbonate anions (HCO_3_^−^), carbonate anions (CO_3_^2−^), protons (H^+^), hydroxide anions (OH^−^), sodium cations (Na^+^), and nitrate anions (NO_3_^−^). The biocathode is treated as a porous electrode with microbial cells of 1-μm diameter and a porosity equivalent to that of a square lattice of spheres (Fig. 1b). We consider the physiology of direct electron transfer through the MtrCAB electron conduit (Fig. 1c) to determine the stoichiometry of CO_2_ reduction to pyruvate for four major CFPs (Fig. 1d-g) using either aerobic or anaerobic nitrate respiration (Fig. 1c).

**Figure 1.**
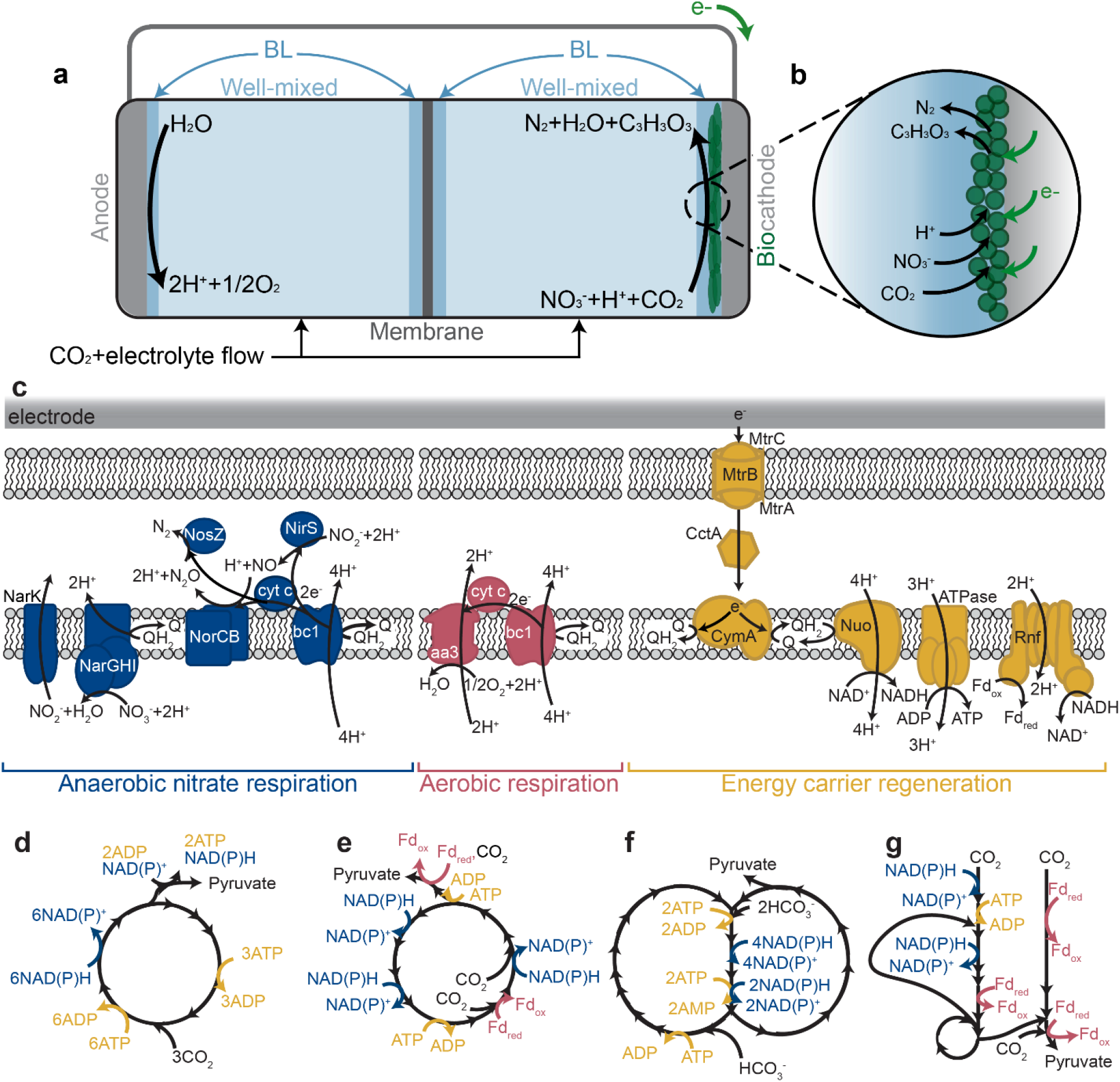
Schematic of a one-dimensional microbial electrosynthesis reactor and direct electron transfer mechanisms for (an)aerobic carbon fixation. (**a**) Reactor scheme. Carbon dioxide (CO_2_) and electrolyte media are fed into well-mixed regions separated by a membrane. (**b**) Direct electron transfer to a microbial biofilm supports carbon fixation to pyruvate using 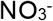 as the terminal electron acceptor. (**c**) Respiratory and energy carrier regeneration mechanisms using 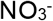 and O_2_ as terminal electron acceptors. (**d**) The Calvin-Benson-Bassham (CBB), (**e**) reductive tricarboxylic acid (rTCA), (**f**) 3-hydroxypropionate bi-cycle, and (**g**) Wood-Ljungdahl pathways for carbon fixation to pyruvate shown with reducing equivalent consumption.

### Comparing microbial respiration and carbon fixation pathways

The terminal electron acceptor and CFP constrain the efficiency and productivity of MES systems. Because O_2_ is more electronegative than NO_3_^−^, aerobic respiration allows microbes to divert a higher fraction of electrons to energy carrier regeneration and therefore carbon fixation (Tables S1 and S2). For microbes using the Calvin-Benson-Bassham (CBB) cycle, aerobic respiration uses only 23.67 electrons per pyruvate molecule, while anaerobic nitrate respiration requires 61.25 electrons for the equivalent reaction. Under anaerobic conditions, obligately anaerobic CFPs use electrons more efficiently than aero-tolerant pathways: the reductive tricarboxylic acid (rTCA) cycle and the Wood-Ljungdahl pathway (WLP) require 47.5 and 46.25 electrons per pyruvate, respectively, while the CBB cycle and the 3-hydroxypropionate (3-HP) bi-cycle need 61.25 and 66.25 electrons. The rate-limiting step in carbon-fixing reactions must also be considered when evaluating productivity. While the WLP uses electrons most efficiently, the enzymatic processes are rate-limited compared to other CFPs (Table S3, Supplementary note 1), so more biomass would be needed to fix carbon at equal rates.

To determine the impacts these competing constraints have on MES systems, we calculated the pyruvate production rate as a function of O_2_ partial pressure for microbes using the CBB cycle to fix carbon with O_2_ as the terminal electron acceptor (Fig. 2a) and pyruvate production versus applied voltage for microbes using NO_3_^−^ as the terminal electron acceptor and different CFPs (Fig. 2b). For aerobic respiration, pyruvate production remains <2 μmol/cm^2^/hr even at O_2_ partial pressures 5-fold greater than in the atmosphere 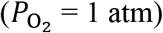 due mainly to the low solubility and corresponding transport limitations of O_2_ in aqueous solutions (Supplementary note 2). In contrast, pyruvate production using anaerobic nitrate respiration reaches ~16.9 μmol/cm^2^/hr at ~2.3 V for microbes using the rTCA cycle before the system becomes CO_2_ transport limited, defined as the point at which the CO_2_ concentration reaches ~0 mM at the base of the biofilm (current collector). Maximum pyruvate production rates for microbes using the CBB cycle (~7.0 μmol/cm^2^/hr), 3-HP cycle (~4.4 μmol/cm^2^/hr), and WLP (~1.9 μmol/cm^2^/hr) are limited by the biomass available to fix carbon in 50 μm biofilms well before CO_2_ transport becomes rate limiting.

**Figure 2.**
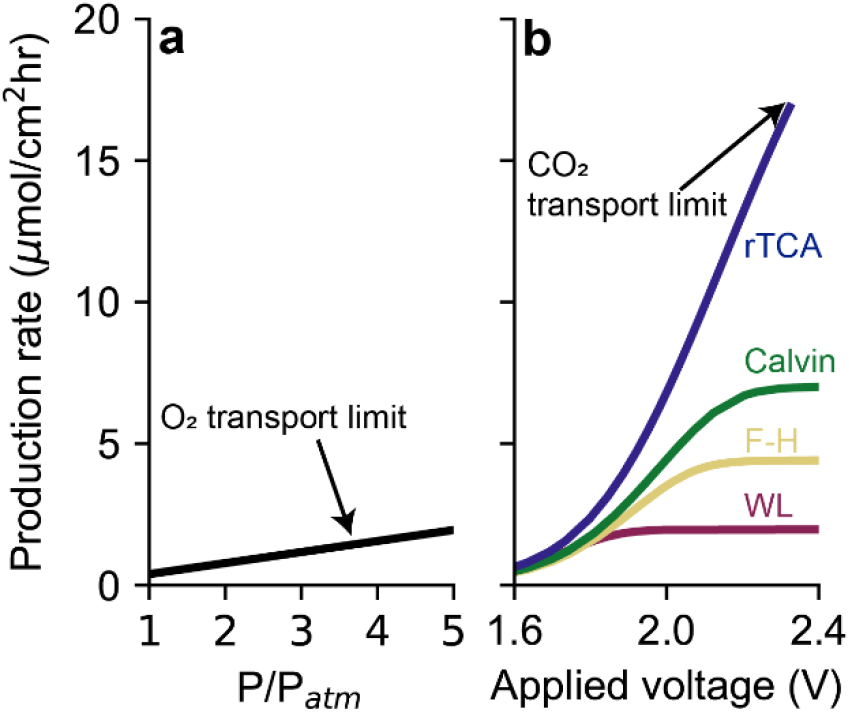
Effects of terminal electron acceptor and carbon fixation pathway on reactor operation. Pyruvate production rate at equivalent biofilm thickness (50 μm) as a function of (**a**) O2 pressure supplied to the reactor headspace relative to atmospheric O_2_ for microbes using the Calvin cycle with O_2_ as the terminal electron acceptor, and (**b**) applied voltage for each carbon fixation pathway with 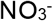 as the terminal electron acceptor. Reactor conditions: initial pH=7.4, 0.25 M NaNO_3_, D=5 hr^−1^.

### Biofilm thickness effects on productivity

The biofilm thickness plays an important and complex role in determining both the total carbon fixation rate and the energy efficiency of an MES system. Increasing the biofilm thickness increases the biomass available to fix carbon, enabling a higher total electron uptake rate by increasing the biomass-limited production rate. However, CO_2_ transport through the biofilm will eventually impose an upper bound on the reaction rate. We plot the voltage necessary to achieve selected pyruvate production rates as a function of biofilm thickness for microbes using the rTCA cycle (Fig. 3a) or CBB cycle (Fig. 3b). For microbes using the CBB cycle, ~3.5-fold thicker biofilms are needed to achieve equivalent production rates because of the lower turnover number for the rate-limiting enzyme, RuBisCo, and the ~29% less efficient use of electrons. These factors also limit the achievable productivity of microbes using the CBB cycle to <12 μmol/cm^2^/hr because thicker biofilms present a longer distance for CO_2_ diffusion.

**Figure 3.**
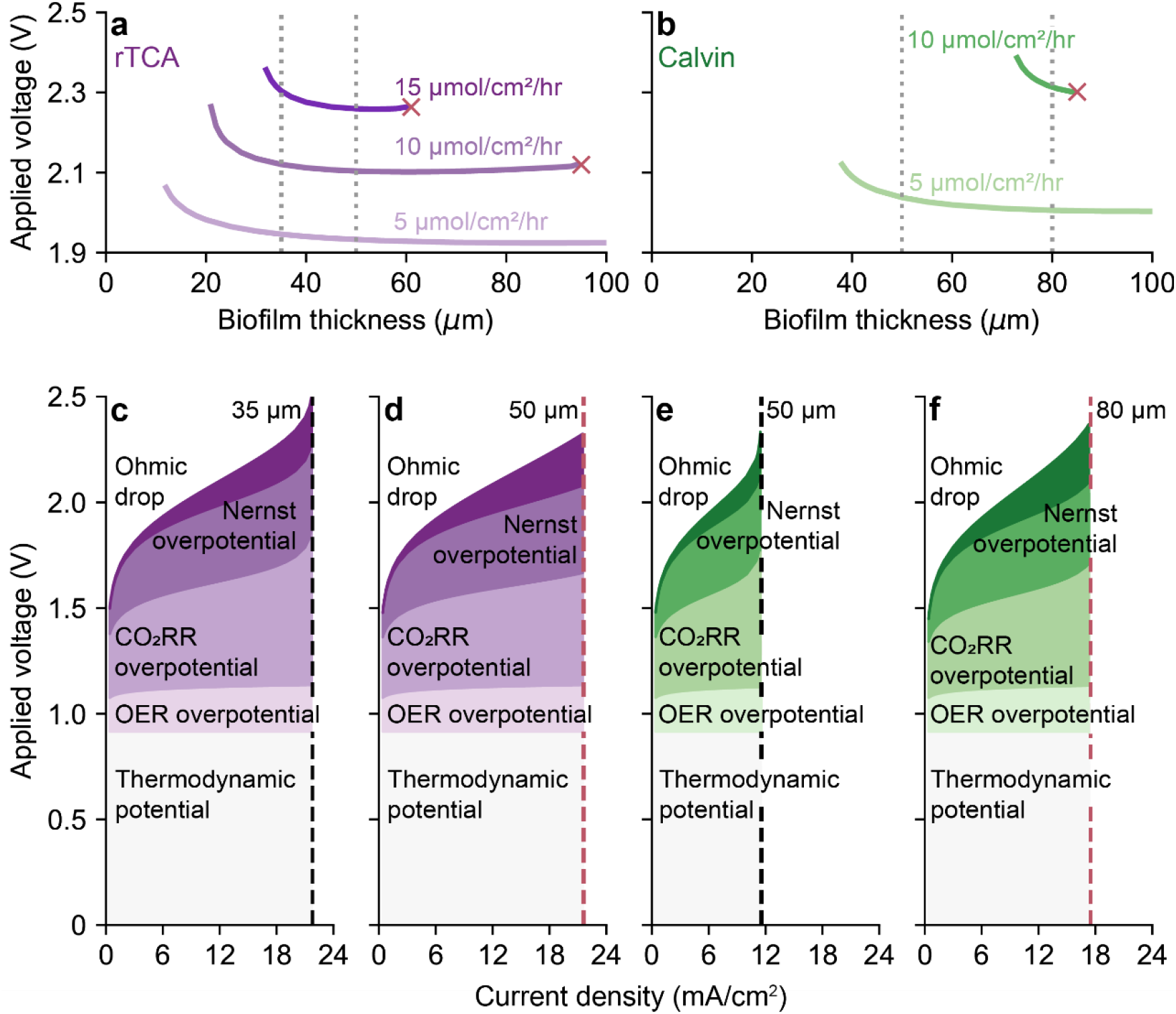
Effect of biofilm thickness on reactor operation. Applied voltage necessary to achieve a specific pyruvate production rate as a function of biofilm thickness for microbes fixing carbon using (**a**) the rTCA cycle and (**b**) the Calvin cycle. Applied voltage breakdown for (**c**) 35 μm, (**d**) 50 μm, (**e**) 50 μm, (**f**) 80 μm biofilms for microbes using the rTCA cycle (**c**, **d**) or Calvin cycle (**e**, **f**). Gray dotted lines in (**a**) and (**b**) correspond to biofilm thicknesses in (**c**–**f**) and were chosen to be representative of different production limits (biomass, CO2 transport). Red crosses in (**a**, **b**) correspond to the CO2 transport limit. Black dashed lines in (**c**, **e**) correspond to biomass-limited current density; red dashed lines in (**d**, **f**) correspond to the CO2 transport-limited current density. Reactor conditions: initial pH=7.4, 0.25 M NaNO_3_, D=5 hr^−1^.

For both CFPs, increasing the biofilm thickness has a non-linear effect on the applied voltage necessary to achieve a fixed production rate (Fig. 3a, b). The initial rapid decline, and the following plateau over a wide thickness range, is due to the competing impacts of the activation overpotential for the CO_2_ reduction reaction (CO_2_RR) and transport-associated (Nernst and Ohmic) overpotentials (Supplementary note 3). To describe these trends, we show the applied voltage breakdown for microbes using the rTCA (Fig. 3c, d) or CBB (Fig. 3e, f) cycles at representative biofilm thicknesses. For microbes using the rTCA cycle, increasing the biofilm thickness from 35 μm (Fig. 3c) to 50 μm (Fig. 3d) increases the maximum current density the biofilm can support from ~21.5 mA/cm^2^ to ~31 mA/cm^2^, but CO_2_ transport restricts the current density to 21.5 mA/cm^2^ for the 50-μm biofilm. Increasing the biofilm thickness reduces the applied voltage necessary to achieve a given production rate (Fig. 3a). The difference is slight at lower rates because increased Ohmic loss (due to electron conduction through a thicker biofilm) mostly balances the reduced activation overpotential associated with the CO_2_RR. However, as the production rate approaches the biomass-limited rate for the 35-μm film, the difference increases significantly, reaching ~200 mV at 21.5 mA/cm^2^ (~16.9 μmol/cm^2^/hr) because the activation overpotential for the 35-μm film rises sharply (Fig. 3c). Similar behavior is observed when comparing 50 μm (Fig. 3e) and 80 μm (Fig. 3f) biofilms using the CBB cycle to fix CO_2_.

### Marker gene phylogenetic analysis

Marker gene phylogeny extractions from the Reference Proteomes database resulted in a total of 5329 genomes encoding at least one marker (Fig. 4; Figs. S4-S12; Table S6). Outer membrane cytochromes and their downstream electron transfer components are modular^30,31^. Therefore, although our model focuses on MtrCAB, we extend our genome mining analysis to include other cytochromes capable of DET, including the whole MtrC/OmcA-family along with the biochemically characterized DmsE/MtrA/PioA/MtoA-, ExtA-, and Cyc2-family proteins (Supplementary note 4). We are not aware of a comprehensive multi-heme cytochrome phylogeny that contains all the clades used in this study. Here, we present this phylogeny (Fig. S11), and show the phylogenetic distribution of these cytochromes (Fig. 4).

**Figure 4.**
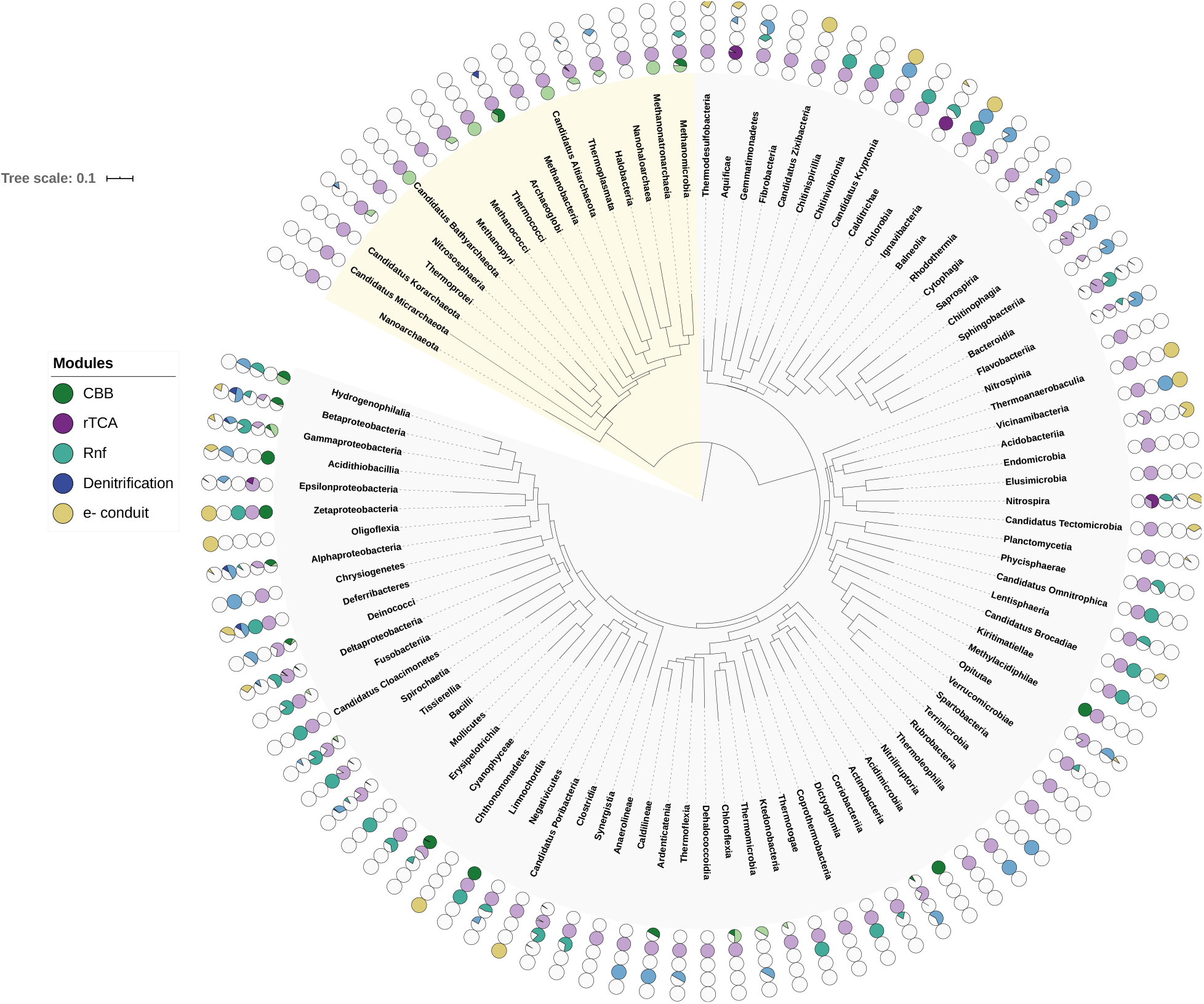
Distribution of target metabolic modules in the Reference Proteomes database. Species were binned by class (or phylum if no class was available), and 16S sequences representing each taxon were used to construct a phylogenetic tree. Each pie chart represents the percentage of species within each taxon that contained the appropriate marker genes. For the CBB (green), rTCA (purple), and denitrification (blue) modules, dark colors represent species that had all marker genes, while lighter colors represent species that had at least one, but not the full set of marker genes. The Rnf (teal) module represents the presence of only one marker gene, while the electron conduit (yellow) represents species that had at least one of the four possible marker genes. Archaea are highlighted in pale yellow, while bacteria are highlighted in pale gray.

We extracted 170 genomes from this dataset that have a complete CBB or rTCA cycle plus at least one electron conduit or that have promise as a chassis for engineering CBB or rTCA-based electrosynthesis (Fig. 5). Of these, 88 organisms have a complete CBB cycle plus at least one electron conduit; genetic engineering methods have been demonstrated in 17 of these organisms. Although 16 organisms have a complete rTCA cycle, only two of these organisms have genetic methods available. The remaining extracted genomes have promise as a chassis for engineering CBB#x002D; or rTCA-based electrosynthesis (Fig. 5). In total, 67 genomes encoded both PFOR and OGOR plus at least one electron conduit but were missing AclB/CcsA. Several genomes where completion of the CBB cycle or addition of an electron conduit is feasible were also included.

**Figure 5.**
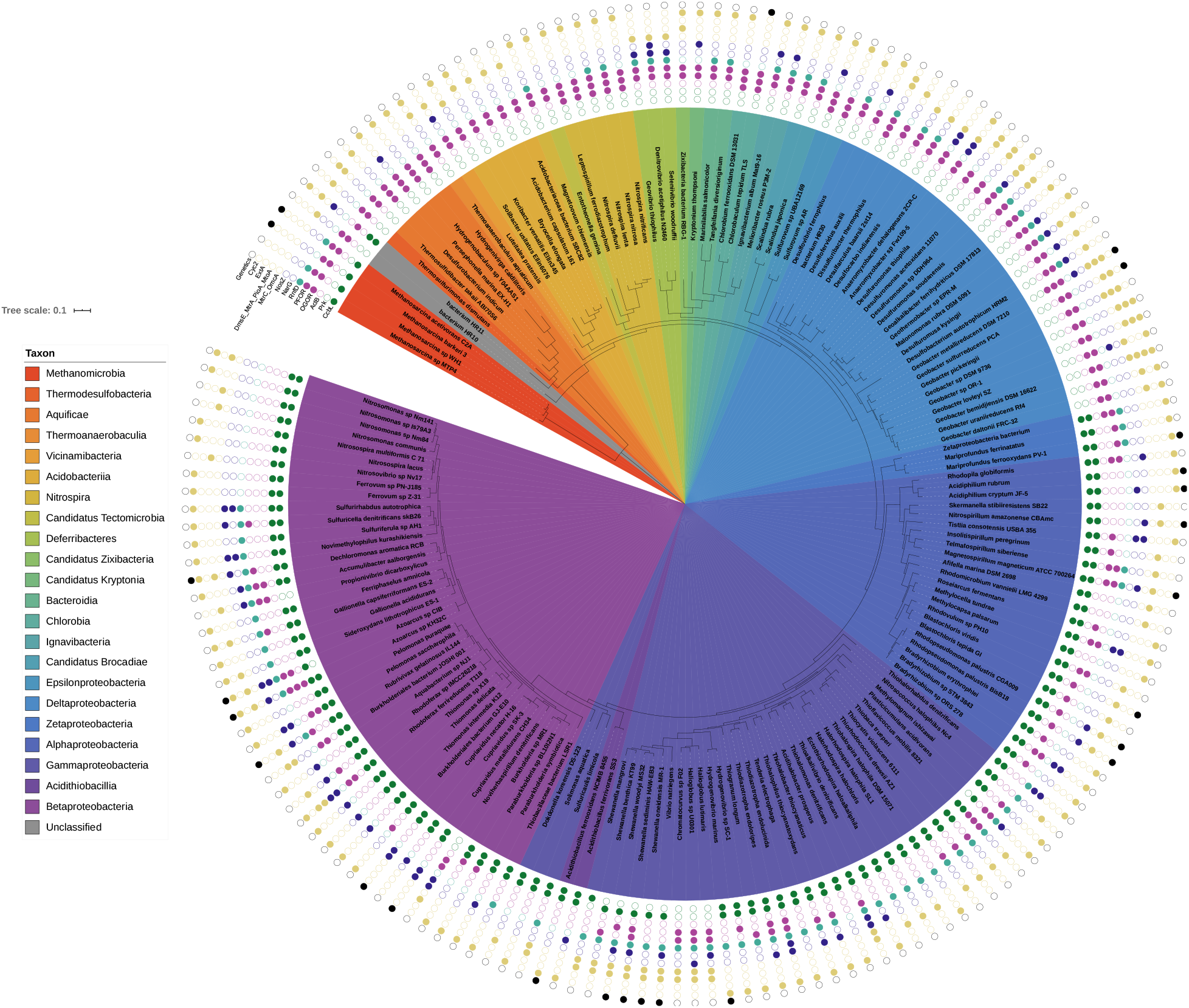
Species that are capable of electroautotrophy or that are potential chassis for engineering electroautotrophy. Marker gene distribution for the CBB (green) cycle, the rTCA (purple) cycle, Rnf (teal), denitrification (blue), and electron conduits (yellow) are displayed on a species-level 16S phylogenetic tree. If a given species has been genetically transformed, that species is marked as having genetics (black).

## Discussion

This analysis has significant implications for MES systems. We show that although O_2_ is a more efficient terminal electron acceptor than NO_3_^−^, low solubility severely limits the productivity of microbes using aerobic respiration. Under anaerobic nitrate respiration, the CO_2_ transport limit imposes an upper bound on productivity, so increasing biofilm thicknesses cannot enable arbitrarily high production rates. For microbes with a lower enzymatic reaction rate limit (lower turnover number), the CO_2_ transport limit is also lower, so microbes using the 3-HP bi-cycle and WLP cannot match the productivity achievable by microbes using the rTCA or CBB cycle regardless of the biofilm thickness. A lower turnover number for the rate-limiting enzyme also increases transport- or activation-associated overpotentials for a given CO_2_-fixation rate, reducing energy efficiency. Combined, these results indicate that microbes using the rTCA cycle are likely to be both the most productive and most efficient biocatalysts for MES systems. Microbes that use the CBB cycle are the second-best option because, although the cycle’s electron utilization efficiency is lower than that for the WLP, the biomass-limited reaction rate is much higher, so the CO_2_ transport-limited production rate is higher and transport-associated inefficiencies are lower.

We therefore used a marker protein phylogeny-driven bioinformatics approach to identify organisms capable of electroautotrophy by coupling electron uptake to nitrate respiration and using either the CBB or rTCA cycles to fix carbon. In our dataset, 19 organisms have complete CBB cycles, NarG, and at least one electron conduit (Fig. 5). To our knowledge, none of these organisms have previously been identified as electroautotrophs and physiological confirmation plus development of genetic tools would expand the available CBB-based electroautotrophs for industrial applications. In contrast, only *Geobacter metallireducens* encodes the rTCA cycle, NarG, and at least one electron conduit. As genetic tools have been developed for *G. metallireducens*^32^, this organism represents an especially promising catalyst for industrial MES. Because we identified only a small number of organisms that have all the desired modules, most of which do not have genetic tools, we also identified organisms that have potential as synthetic chassis for MES, which we discuss here in the context of several possible synthetic biology strategies for engineering electroautotrophy.

First, an organism may have a partial CFP that can be completed by heterologous expression of the missing components. Several recent demonstrations make this an attractive strategy: the CBB cycle has been engineered into heterotrophs by the addition of key enzymes^33–35^, and the rTCA cycle was completed in *Geobacter sulfurreducens* to enable electroautotrophy using both rational engineering^36^ and directed evolution strategies^37^. An alternate CFP, the reductive glycine pathway^38,39^, is a third option since its modularity has recently been confirmed by functional expression in both *Escherichia coli*^40^ and *Cupriavidus necator*^41^. Our dataset revealed several organisms in which completion of a partial CFP may be viable. Four of five *Shewanella* species identified in this study encode Prk, three encode partial rTCA cycle markers, and two encode NarG. Given the prevalence of genetic tools available to this genus, *Shewanella oneidensis*’ well-characterized use in bioelectrochemical systems, and facultative anaerobic metabolism, engineering a complete CFP in *Shewanella* could lead to readily-engineerable electroautotrophs. All *Shewanella* species encode the Rnf marker gene, indicating that they can support the efficient reverse-electron transport-driven ferredoxin reduction that may be required for the rTCA cycle.

Second, an organism encoding a complete CFP may be engineered to directly uptake electrons from a cathode. The electron conduit native to *Shewanella oneidensis*, MtrCAB/CymA, has been functionally expressed in *E. coli*^42–44^, which is a promising host for this strategy since multiple CFPs have been successfully engineered and it naturally respires nitrate. Alternatively, *Cupriavidus spp.* encode both a complete CBB cycle and nitrate respiration and have genetic tools available^45^, making these species an attractive option for expressing an electron conduit. Because only two organisms that encode a full rTCA cycle with at least one electron conduit have genetic methods available, the green sulfur bacterium *Chlorobaculum tepidum*, which encodes the full rTCA and has genetic tools, may be another suitable host for electron conduit expression.

Finally, an organism with a complete CFP and a functional electron conduit may be engineered to use an alternate electron acceptor. We use nitrate respiration in our model since it is the most thermodynamically favorable soluble electron acceptor; nitrate respiration could be introduced in organisms by expressing the NarGHI complex. However, 5 molybdopterin cofactor biosynthesis enzymes are also necessary for proper functioning of this complex, so an appropriate chassis would benefit significantly from natural expression of these supporting proteins. *Rhodopseudomonas palustris* is an excellent candidate for this strategy since it has a characterized electron conduit^23^, a complete CBB cycle, and encodes the molybdopterin biosynthesis genes. This organism has been engineered for *poly*-hydroxybutyrate^46^ and *n*-butanol^47^ production in MES systems, so heterologous expression of NarGHI may enable higher yields and productivities. *Azoarcus* sp. KH32C has complete CBB and rTCA cycles and encodes NosZ, while lacking NarG. A genetic system was developed in the related *Azoarcus* sp. strain BH72^48^, which may open up KH32C for heterologous expression of NarGHI, further increasing the potential of this species for MES.

The multiheme cytochrome phylogeny developed here (Fig. S11) indicates that a significant number of undiscovered cytochromes that support electron exchange with an electrode may exist. This hypothesis is supported by reports of direct electron uptake independent of the four biochemically characterized families of outer membrane cytochromes used in our study: both the methanogenic archaeon *Methanosarcina barkeri*^49^ and the green sulfur bacterium *Prosthecochloris aestuarii*^50^ have been shown to directly accept electrons from a cathode. Cytochromes involved in direct electron transfer may therefore be more widespread and diverse than currently realized, opening the possibility of a significant number of additional chassis for MES (Supplementary note 4).

Here we have identified several microbial chassis that have potential as industrial MES strains (Table 1) and outlined a series of Technology Readiness Levels (TRLs) to evaluate industrial relevance of microbial catalysts for MES (Table S7). Beyond the genetic modules that enable electroautotrophy, additional factors constrain the productivity achievable by a given organism. For current densities >10 mA/cm^2^, which are likely necessary for viable production capacity, our model predicts alkaline conditions throughout the biofilm, indicating that (facultative) alkalophilicity is a desirable trait in the ideal MES strain. A higher salinity reduces Ohmic overpotential and an increased bicarbonate concentration can enhance productivity by aiding CO_2_ transport, so halophilicity or halotolerance is similarly advantageous. Unfortunately, the pH- and halo-tolerance of the organisms we identify is unclear, so future studies, in addition to confirming electroautotrophic capacity, should also characterize these traits. A suitable microbial catalyst could also be engineered to tolerate alkaline or saline conditions using rational engineering or directed evolution strategies. Because the turnover number of the rate-limiting enzyme plays a key role in setting productivity and efficiency limits for MES, these values should also be characterized in the organism(s) of interest. Strain engineering can then focus on increasing the rate limit either by increasing the enzyme turnover number or overexpressing the rate-limiting enzyme.

**Table 1.**
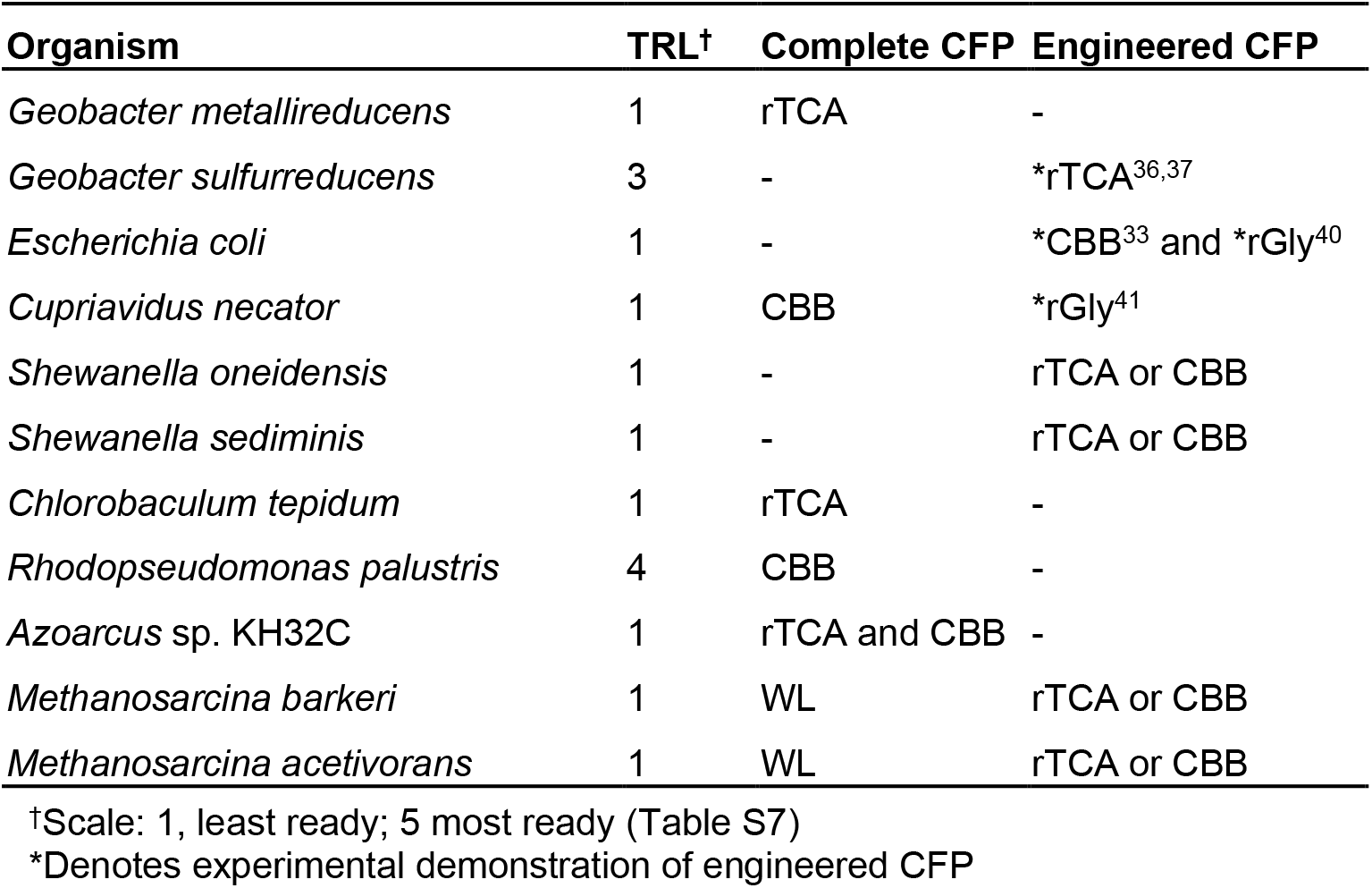
Promising microbial chassis for MES.

MES strain selection and engineering should also be guided by the desired product. For example, the CBB cycle is well-suited for the production of sugars such as sucrose because its end product is glyceraldehyde-3-phosphate, while the rTCA cycle may be better suited to fatty acid production since the end product, acetyl-CoA, is used directly by fatty acid biosynthesis pathways. Coupling direct electron uptake-driven carbon fixation to the biosynthesis of platform chemicals will enable a sustainable process for the conversion of CO_2_ into renewable commodities.

## Acknowledgements

This work was supported by the Center for the Utilization of Biological Engineering in Space (CUBES, https://cubes.space), a NASA Space Technology Research Institute (grant number NNX17AJ31G). A.J.A. is supported by an NSF Graduate Research Fellowship under grant number DGE 1752814. We thank Dr. Lien-Chun Weng (UC Berkeley) for advice on COMSOL modeling, Helen Bergstrom (UC Berkeley) for useful discussions on electrochemistry, and Dr. Paul Tol (Netherlands Institute for Space Research, SRON) for a helpful reference on accessible color schemes (https://personal.sron.nl/~pault/).

## Author contributions

A.J.A. conceived of the idea, developed the model, and analyzed data. J.M.H. performed the phylogenetic analysis and analyzed data. A.J.A. and J. M. H. wrote the manuscript. A.P.A and D.S.C. edited the manuscript and supervised the project.

## Competing interests

The authors have no competing interests to declare.

## Data availability

Data and code used to generate figures are available at 10.6084/m9.figshare.13224617.

## Computational Methods

### System overview

Species transport for an open electrochemical system must satisfy mass conservation:

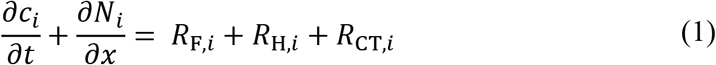

where *c*_*i*_ is the concentration, *N*_*i*_ is the molar flux, and *R*_F,*i*_, *R*_H,*i*_, and *R*_CT,*i*_ are the net volumetric rates of formation or consumption for species *i* (CO_2_, HCO_3_^−^, CO_3_ ^2−^, H^+^, OH^−^, Na^+^, NO_3_^−^) due to gas and electrolyte (F) feed terms, homogeneous (H) chemical reactions, and electrochemical charge transfer (CT) reactions, respectively. *R*_F,*i*_ applies only in the well-mixed electrolyte phases where gas and electrolyte feeds are introduced and *R*_CT,*i*_ applies only in the porous biocathode layer.

In the following sections, we formulate the equations that govern transport and reactions within the MES system, describe assumptions, and report the key parameter values used in our model.

### Species transport in the electrolyte boundary layers, membrane, and porous biocathode

The molar flux of species (assuming no net fluid velocity) in dilute electrolyte solutions is written as the sum of diffusive and migrative fluxes:

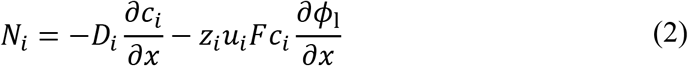

where *D*_*i*_ and *u*_*i*_ are the diffusivity and mobility (related by the Nernst-Einstein relationship, *u*_*i*_ = *D*_*i*_/*RT* for dilute solutions) of species *i*, *z*_*i*_ is the charge number, *F* is Faraday’s constant, and *ϕ*_*l*_ is the local electrolyte potential. In the anion exchange membrane (AEM), we reduce diffusion coefficients of anions and cations by a factor of 10 and 100 respectively relative to those in the electrolyte to model a generic anion exchange membrane and assume a fixed background positive unit charge with a 0.5 M concentration, following Singh *et al.*^1^ We also use effective diffusion coefficients within the biocathode layer calculated using the Bruggeman relationship,

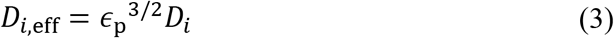

where *∊*_p_ is the biofilm porosity. The net ionic current density in the electrolyte (*i*_l_) can be calculated from the total ionic flux:

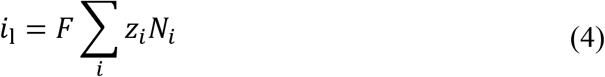

where *F* is Faraday’s constant and *z*_*i*_ is the charge number, following electroneutrality,

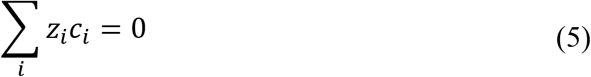

### Gas feed and electrolyte flow in the well-mixed electrolyte

The well-mixed electrolyte regions are assumed to have sufficient convective mixing such that no concentration gradients are formed. Species transport into and out of the boundary layers is considered at the interface between the well-mixed and boundary layer electrolyte phases (Fig. 1a, b). Constant gas feed and electrolyte flow terms in the well-mixed regions are included to describe a continuously operating system, given by:

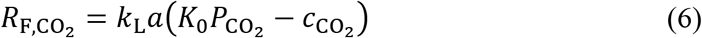

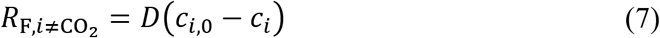

where *k*_L_*a* is the volumetric mass-transfer coefficient (in units s^−1^) on the liquid side of the gas/ liquid interface, *K*_0_ is Henry’s constant for CO_2_ in water, 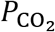 is the pressure of CO_2_ in the gas phase, *D* is the dilution rate (defined as the inverse space time, or volumetric flow rate divided by reactor volume), and *c*_*i*,0_ is the initial or feed concentration of the *i*th species. The equilibrium of CO_2_ between the gas and liquid phases, 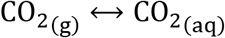, is describedby Henry’s constant such that:

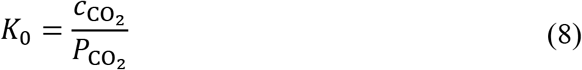

Henry’s constant for CO_2_ depends on the temperature and salinity of the aqueous phase and follows an empirical relationship^2^,

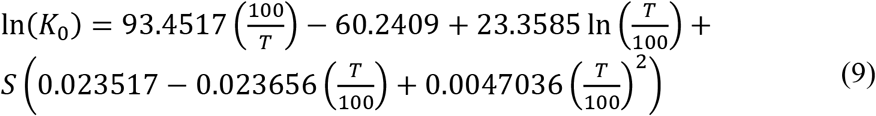

where *S* is the salinity in units g/kg and *T* is the temperature.

### Homogeneous chemical reactions

The acid-base bicarbonate/carbonate and water-dissociation reactions shown below occur in all phases and are treated as kinetic expressions without assuming equilibrium (eq. 15):

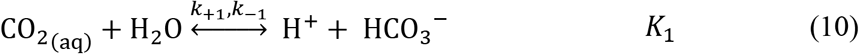

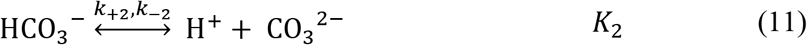

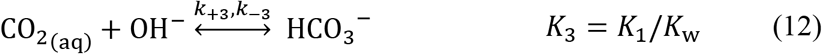

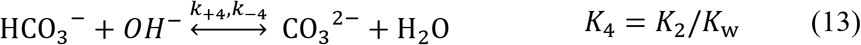

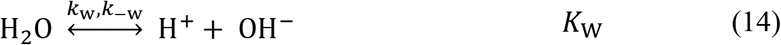

where *k*_+*n*_ and *k*_−*n*_ are the forward and reverse rate constants, respectively, and *K*_*n*_ is the equilibrium constant for the *n*th reaction. Source and sink terms resulting from these reactions are compiled in *R*_H,*i*_, written as:

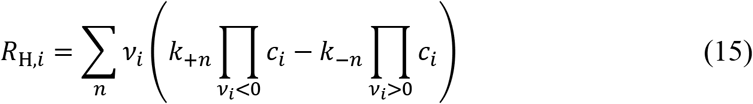

where *v*_*i*_ is the stoichiometric coefficient of species *i* for the *n*th reaction and reverse rate constants are calculated from:

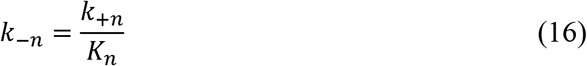

### Electrode reactions – anode

The surface reaction at the anode is the oxidation of water:

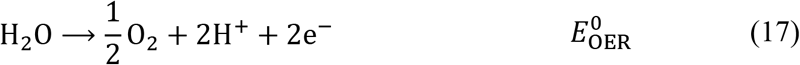

where 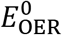 is the equilibrium potential of the oxygen evolution half-cell reaction (OER) at standard state. The anode reaction is related to species transport by a flux boundary condition at the electrode surface,

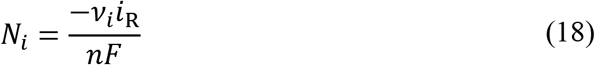

where *i*_R_ is the reaction current density and *n* is the number of electrons participating in the electrode reaction. We model charge transfer kinetics at the anode using Butler-Volmer kinetics:

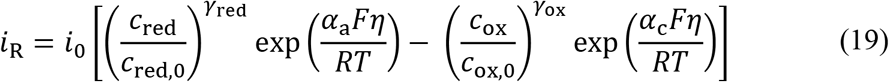

where *i*_0_ is the constant exchange current density, *γ*_red/ox_ is the reaction order with respect to a reactant, *α*_a/c_ is the anodic/cathodic transfer coefficient, and *η* is the overpotential. The overpotential is defined according to:

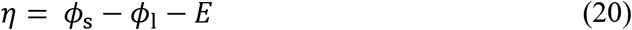

where *ϕ*_s_ is the electrode potential, *ϕ*_l_ is the electrolyte potential, and *E* is the half-cell equilibrium potential.

Because water oxidation creates acidic conditions near the anode surface, bicarbonate and carbonate species will be converted to aqueous CO_2_ according to Le Chatelier’s principle. To avoid the unrealistic supersaturation of CO_2_ in the electrolyte this would cause, we describe evolution of CO_2_ in the electrolyte (Supplementary note 6) as:

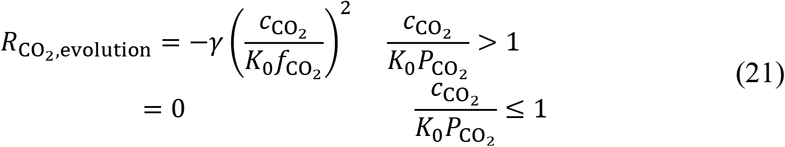

where *γ* is the releasing coefficient and 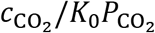 is the supersaturation ratio. This formulation was originally reported by Wilt, and was utilized to describe CO_2_ evolution in abiotic electrochemical systems previously^1,3–5^.

### Electrode reactions – biocathode

We consider the physiology of direct electron transfer through the MtrCAB electron conduit (Fig. 1c in the main text) to determine the stoichiometry of CO_2_ reduction to pyruvate for four major carbon fixation pathways (Fig. 1d–g in the main text) using either aerobic or anaerobic nitrate respiration. All processes start with quinone (Q) reduction,

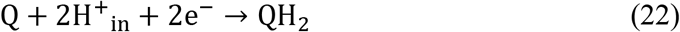

using the MtrCAB/CctA/CymA electron conduit native to *S. oneidensis*^6–9^. For aerobic respiration, the respiratory complex III (*e.g.* the *bc1* complex) oxidizes a quinol, pumping protons across the inner membrane^10,11^:

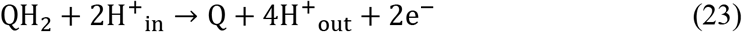

where the subscripts “in” and “out” refer to ion locations in the intracellular space and periplasm, respectively. The two electrons liberated in this process are transported by *c*-type cytochromes to respiratory complex IV (*e.g.* the *aa3* complex), which transports two additional protons across the inner membrane and reduces O_2_ to H_2_O^10,12^:

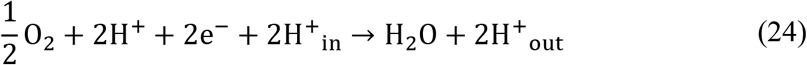

For anaerobic nitrate respiration, quinols are consumed both to pump protons via respiratory complex III, eq. (23), and to reduce NO_3_^−^ to nitrite (NO_2_^-^) using, *e.g.* the Nar complex^13^:

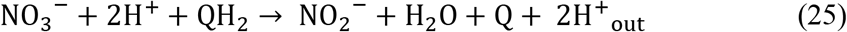

Further reactions consume electrons liberated by quinol oxidation to complete the reduction of NO_2_^−^to N_2_^13^:

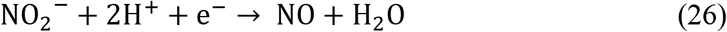

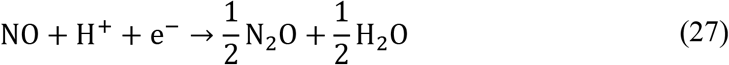

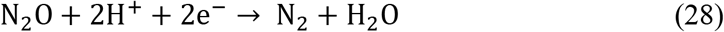

Carbon fixation pathways require NAD(P)H, ATP, and/or reduced ferredoxins (Fd_red_) as reducing equivalents^14^. Cells can regenerate these reducing equivalents by translocating protons (PMF consumption) using, *e.g.*, the Nuo complex for NADH^15^, ATP synthase for ATP^16^, and the Rnf complex for ferredoxins^17–19^ according to:

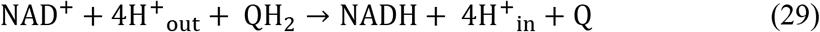

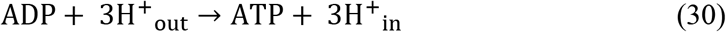

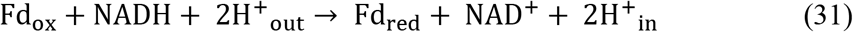

We use these regeneration mechanisms to determine the stoichiometry (number of reduced molecules produced per number of electrons consumed) for aerobic or anaerobic nitrate respiration (Table S1). Because carbon fixation pathways have different energy carrier requirements, we also derive the overall stoichiometry for CO_2_ reduction to pyruvic acid (pyruvate) (Table S2). For the aero-tolerant carbon fixation pathways (Calvin cycle, eq. (32), Fuchs-Holo bi-cycle, eq. (33)), the cathodic half-cell reactions using aerobic respiration are:

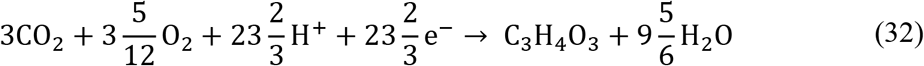

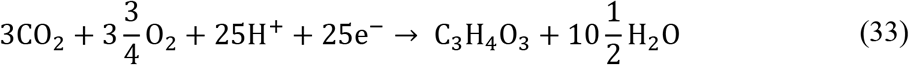

For carbon fixation pathways using NO_3_^−^ as the terminal electron acceptor, the half-cell reactions are:

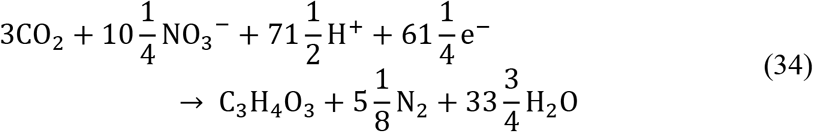

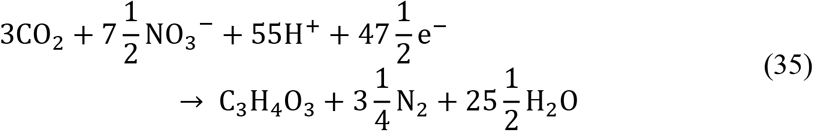

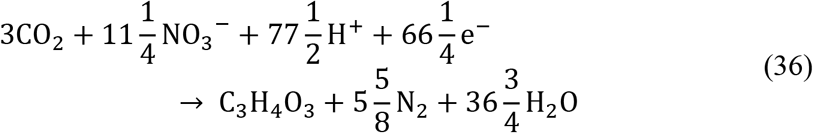

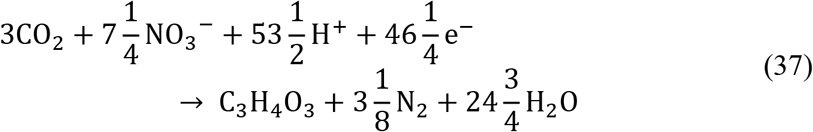

where eq. (34) is for the Calvin cycle, eq. (35) is for the rTCA cycle, eq. (36) is for the Fuchs-Holo bicycle (F-H), and eq. (37) is for the Wood-Ljungdahl (WL) pathway.

Biocathode reactions, eq. (32–37), relate CO_2_-fixing reactions to species transport in the biocathode layer by,

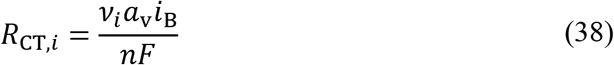

where *a*_v_ is the active specific surface area of the biocathode and *i*_B_ is the current density on the biocathode surfaces. The active specific surface area is calculated based on the geometric assumptions described above, resulting in:

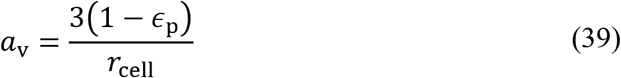

where *r*_cell_ is the radius of the spherical microbe. The current density can be limited by the maximum rate at which the cells can fix CO_2_, which depends on the turnover number of the rate-limiting enzyme in the carbon fixation pathway. To account for this, we impose a limit on *i*_B_ via:

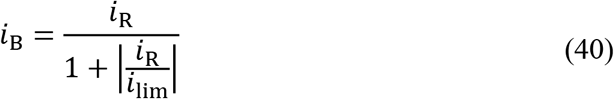

where *i*_lim_ is the biomass-limited current density. We calculate the biomass-limited current density by projecting the enzymatic rate limit to the total cell surface:

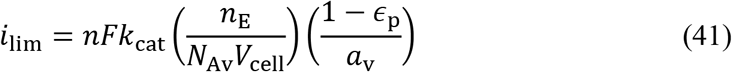

where *k*_cat_ is the enzyme turnover number (units s^−1^), *n*_E_ is the enzyme amount in each cell (units cell^−1^) *N*_Av_ is Avogadro’s number, and *V*_cell_ is the microbe volume. This formulation for the limiting current density relies on the fact that the rate of intracellular diffusion of substrates is much faster than the rate-limiting reaction step in carbon fixation pathways (see supplementary note 5 for calculations), indicating that energy carriers and CO_2_ have complete and effectively immediate access to intracellular enzymes once generated at or delivered to the cell surface. We model charge transfer using Butler-Volmer kinetics, eq. (19).

### Electron transport in the solid electrode

Electron transport in the solid electrode regions is governed by charge conservation and Ohm’s law, given by:

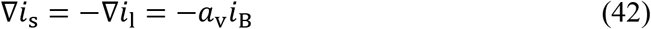

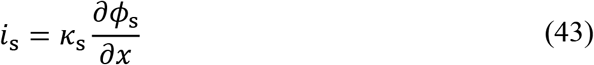

where *i*_s_ is the electrode current density and *κ*_s_ is the anode/biocathode conductivity. The conductivity in the biocathode is modified by a Bruggeman correction:

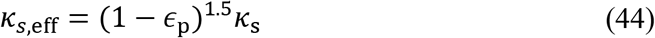

### Numerical method

The governing equations are solved using the MUMPS general solver in COMSOL Multiphysics 5.4. The modeling domain has a maximum element size of 10 μm in the well-mixed regions and 0.5 μm near boundaries to capture steep concentration gradients; the solution was independent of increasing mesh resolution. Model parameters are listed in Table 3. The potential in the reactor is calculated relative to zero potential at the cathode base and potential or current density is applied as a boundary condition at the anode.

### Construction and analysis of phylogenetic trees

Seed sequences for marker genes^20–22^ of each pathway (Table S4) were fed to JackHMMER for homology searches^23^. Protein identifiers from the JackHMMER output were used to retrieve full-length protein sequences from Uniprot (The Uniprot Consortium 2019). Protein alignments were created using MAFTT^24,25^. Maximum likelihood trees for each protein search result were built using FastTree 2^26^. Trees were visualized with Iroki^27^. Trees were manually annotated through a combination of conserved protein domain information (The Uniprot Consortium 2019), inclusion of characterized proteins when available^28,29^, and nearest characterized protein information for uncharacterized proteins^29^. Protein sequences corresponding to each marker gene were extracted from the phylogenies using the R package ape^30^. Extracted protein sequences for all marker genes were used to construct a table containing gene presence or absence information for each genome in the dataset (Table S6). Phylogenetic trees displaying gene presence and absence data were built using 16S sequences that were aligned using MAFFT^24,25^, built using FastTree 2^26^, and visualized using the Interactive Tree of Life^31^ and annotated using taxonomy data from NCBI. Presence of genetic tools was established through a literature search (Table S5).

MtrC from *Shewanella oneidensis* was used as the seed sequence in generating the multi-heme cytochrome phylogeny (Table S4). The JackHMMER output was filtered to exclude sequences with less than eight heme motifs (CxxCH or CxxxCH), and those sequences outside the range of sequence lengths of sequences containing exactly ten heme motifs (188-1434 amino acids). Biochemically characterized multi-heme cytochromes were used to annotate the phylogeny: MtrC^32^, OmcA^33^, MtrF^33^, PioA^34,35^, MtoA^36^, DmsE^37^, MtrA^33^, MtrD^33^, and ExtA^38^. Although MtrA/D are components of three-subunit electron conduits with MtrC/F, members of this clade, such as PioA and MtoA, are components of two-subunit electron conduits. While DmsE is characterized as being a DMSO reductase, this protein clustered very closely to MtrA and MtrD, and we were therefore unable to separate putative DMSO reductases from cytochromes that may interact with an electrode. Thus, all members of the DmsE/MtrA/PioA/MtoA-family clade of proteins were kept for downstream analysis as potential multi-heme cytochromes involved in direct electron transfer with an electrode.

## Notes

https://doi.org/10.6084/m9.figshare.13224617

### Competing Interest Statement

The authors have declared no competing interest.

