## Supplementary information for "Systems-informed genome mining for electroautotrophic microbial production"

**Affiliations**
1. Department of Chemical and Biomolecular Engineering, University of California, Berkeley, CA 94720, USA
2. Department of Bioengineering, University of California, Berkeley, CA 94720, USA
3. Environmental Genomics and Systems Biology Division, Lawrence Berkeley National Laboratory, 1 Cyclotron Road, Berkeley, CA 94720, USA
4. Molecular Biophysics and Integrated Bioimaging Division, Lawrence Berkeley National Laboratory, 1 Cyclotron Road, Berkeley, CA 94720, USA

[†]These authors contributed equally.

### Supplementary Tables

**Supplementary Table 1. Stoichiometric analysis for energy carrier regeneration with $O_2$ and $NO_3^-$ respiration.**

| Terminal electron acceptor | Proton motive force ($H^+$ to periplasm) | $QH_2$ | NADH | ATP | $Fd_{red}$ |
|---|---|---|---|---|---|
| $O_2$ | $3/e^-$ | $1/2e^-$ | $1/3.33e^-$ | $1/e^-$ | $1/4e^-$ |
| $NO_3^-$ | $0.8/e^-$ | $1/2e^-$ | $1/7e^-$ | $1/3.75e^-$ | $1/9.5e^-$ |

**Supplementary Table 2. Energetic requirements of carbon fixation for pyruvate production.**

| Pathway | Energetic requirements to fix one mole of pyruvate | | | | |
|---|---|---|---|---|---|
| | Mol NAD(P)H | Mol ATP | Mol $Fd_{red}$ | Mol $e^-$ | Reference |
| Calvin ($O_2$) | 5 | 7 | 0 | 23.67 | [1] |
| Calvin ($NO_3^-$) | 5 | 7 | 0 | 61.25 | [1] |
| Reductive TCA | 3 | 2 | 2 | 47.5 | [2] |
| Fuchs-Holo ($O_2$) | 6 | 7 | 0 | 25 | [3,4] |
| Fuchs-Holo ($NO_3^-$) | 6 | 7 | 0 | 66.25 | [3,4] |
| Wood-Ljungdahl | 2 | 1 | 3 | 46.25 | [5] |

**Supplementary Table 3. Model parameters.**

| Parameter | Value | Unit | Reference/Notes |
|---|---|---|---|
| Operating conditions | | | |
| $T$ | 298 | K | |
| $P$ | 1 | atm | |
| $D$ | 1 | $hr^{-1}$ | |
| $pH_0$ | 7.4 | -- | |
| $C_{NaNO_3,0}$ | 0.250 | M | |
| Reactor geometry | | | |
| $l_{bl}$ (boundary layer thickness) | 100 | μm | Supplementary note 7 |
| $l_m$ (membrane thickness) | 100 | μm | Assumed |
| $\epsilon_p$ (biocathode porosity) | 0.48 | -- | Assumed |
| $r_{cell}$ (cell radius) | 0.5 | μm | Assumed |
| Diffusion coefficients | | | |
| $D_{Na^+}$ | $1.34 \times 10^{-5}$ | $cm^2\ s^{-1}$ | [6] |
| $D_{NO_3^-}$ | $1.7 \times 10^{-5}$ | $cm^2\ s^{-1}$ | [7] |
| $D_{H^+}$ | $9.311 \times 10^{-5}$ | $cm^2\ s^{-1}$ | [8] |
| $D_{OH^-}$ | $5.293 \times 10^{-5}$ | $cm^2\ s^{-1}$ | [8] |
| $D_{HCO_3^-}$ | $1.185 \times 10^{-5}$ | $cm^2\ s^{-1}$ | [8] |
| $D_{CO_3^{2-}}$ | $5.293 \times 10^{-5}$ | $cm^2\ s^{-1}$ | [8] |
| $D_{CO_2}$ | $1.91 \times 10^{-5}$ | $cm^2\ s^{-1}$ | [9] |
| Homogeneous reactions | | | |
| $K_1$ | $10^{-6.37}$ | M | [10] |
| $k_{+1}$ | $3.71 \times 10^{-2}$ | $s^{-1}$ | [10] |
| $K_2$ | $10^{-10.32}$ | M | [10] |
| $k_{+2}$ | 59.44 | $s^{-1}$ | [10] |
| $k_{+3}$ | $2.23 \times 10^3$ | $L\ mol^{-1}\ s^{-1}$ | [10] |
| $k_{+4}$ | $6.0 \times 10^9$ | $L\ mol^{-1}\ s^{-1}$ | [10] |
| $K_w$ | $10^{-14}$ | $M^2$ | [11] |
| $k_{+w}$ | $2.4 \times 10^{-5}$ | $mol\ L^{-1}\ s^{-1}$ | [11] |
| Charge transfer reactions | | | |
| $E^0_{OER}$ | 1.23 | V | [11] |
| $i_{0,OER}$ | $1 \times 10^{-8}$ | $A\ cm^{-2}$ | [12] |
| $\alpha_{a,OER}$ | 1.7 | -- | [12] |
| $\alpha_{c,OER}$ | 0.1 | -- | [12] |
| $E^0_{CO_2RR}$ | 0.314 | V | Supplementary note 8 |
| $i_{0,CO_2RR}$ | $1.38 \times 10^{-4}$ | $A\ cm^{-2}$ | [13] |
| $\alpha_{a,CO_2RR}$ | 0.5 | -- | [13] |
| $\alpha_{c,CO_2RR}$ | 0.5 | -- | [13] |
| Enzyme kinetics | | | |
| $k_{cat,CBB}$ | 10 | $s^{-1}$ | Supplementary note 1 |
| $k_{cat,rTCA}$ | 17.5 | $s^{-1}$ | Supplementary note 1 |
| $k_{cat,FH}$ | 2.1 | $s^{-1}$ | Supplementary note 1 |
| $k_{cat,WL}$ | 0.47 | $s^{-1}$ | Supplementary note 1 |
| $n_{E,CBB}$ | $7.08 \times 10^4$ | $cell^{-1}$ | Supplementary note 1 |

| | | | |
|---|---|---|---|
| $n_{\mathrm{E,rTCA}}$ | $7.08 \times 10^4$ | $\mathrm{cell}^{-1}$ | Supplementary note 1 |
| $n_{\mathrm{E,FH}}$ | $7.08 \times 10^4$ | $\mathrm{cell}^{-1}$ | Supplementary note 1 |
| $n_{\mathrm{E,WL}}$ | $2.124 \times 10^5$ | $\mathrm{cell}^{-1}$ | Supplementary note 1 |
| Electrode conductivity | | | |
| $\kappa_{\mathrm{s,anode}}$ | $1 \times 10^4$ | $\mathrm{S\ m^{-1}}$ | Assumed |
| $\kappa_{\mathrm{s,biocathode}}$ | $1 \times 10^{-1}$ | $\mathrm{S\ m^{-1}}$ | Supplementary note 9 |
| Gas-liquid mass transfer | | | |
| $k_{\mathrm{L}}a$ | $5 \times 10^{-1}$ | $\mathrm{s^{-1}}$ | Supplementary note 6 |
| $\gamma$ | $1.25 \times 10^4$ | $\mathrm{mol\ m^{-3}\ s^{-1}}$ | [14] |

**Supplementary Table 4. Marker genes and seed sequences**

| Pathway/Module | Marker gene | Seed sequence(s) |
|---|---|---|
| CBB | CcbL | Q8KC02_CHLTE |
| CBB | Prk | KPPR_SYNY3; KPPR1_RHOSH; KPPR2_CUPNH |
| rTCA | AclB/CcsA | Q9AQH6_CHLLI |
| rTCA | OGOR | A0A1Q2RUM5_HELPX |
| rTCA | PFOR | A0A402E4E5_HELPX; Q8KC02_CHLTE |
| Fd reduction | RnfD | RNFD_CLOLD |
| Denitrification | NarG | NARG_ECOLI |
| Denitrification | NosZ | NOSZ_PSEST |
| e- conduit | MtrC | Q8EG34_SHEON |
| e- conduit | Cyc2 | A0A0H3ZGZ2_9PROT; O33823_ACIFR |

**Supplementary Table 5. References for organisms from Figure 5 with genetic tools**

| Species | Strain | Reference |
|---|---|---|
| *Acidiphilium cryptum* | JF-5 | [15] |
| *Acidiphilium rubrum* | | [15] |
| *Acidithiobacillus ferrooxidans* | ATCC 23270 / DSM 14882 / CIP 104768 / NCIMB 8455 | [16] |
| *Azoarcus sp.* | CIB | [17] |
| *Azoarcus sp.* | KH32C | [17] |
| *Blastochloris viridis* | | [18] |
| *Chlorobaculum tepidum* | ATCC 49652 / DSM 12025 / NBRC 103806 / TLS | [19] |
| *Cupriavidus metallidurans* | ATCC 43123 / DSM 2839 / NBRC 102507 / CH34 | [20] |
| *Cupriavidus necator* | ATCC 17699 / H16 / DSM 428 / Stanier 337 | [21] |
| *Dechloromonas aromatica* | RCB | [22] |
| *Geobacter metallireducens* | GS-15 / ATCC 53774 / DSM 7210 | [23] |
| *Geobacter sulfurreducens* | ATCC 51573 / DSM 12127 / PCA | [24] |
| *Hydrogenovibrio marinus* | | [25] |
| *Magnetospirillum magneticum* | AMB-1 / ATCC 700264 | [26] |

| | | |
|---|---|---|
| *Mariprofundus ferrooxydans* | PV-1 | [27] |
| *Methanosarcina acetivorans* | ATCC 35395 / DSM 2834 / JCM 12185 / C2A | [28] |
| *Methanosarcina barkeri* | 3 | [29] |
| *Nitrospirillum amazonense* | CBAmc | [30] |
| *Rhodopseudomonas palustris* | BisB18 | |
| *Rhodopseudomonas palustris* | ATCC BAA-98 / CGA009 | |
| *Rubrivivax gelatinosus* | NBRC 100245 / IL144 | [31] |
| *Shewanella oneidensis* | MR-1 | [32] |
| *Shewanella sediminis* | HAW-EB3 | [33] |
| *Shewanella woodyi* | ATCC 51908 / MS32 | [34] |
| *Vibrio natriegens* | | [35] |

**Supplementary Table 7. Technology readiness levels (TRLs) for MES**

| TRL | Description |
|---|---|
| 1 | MES capable (predicted) |
| 2 | MES capable (experimentally verified) |
| 3 | MES capable AND genetic tools (single gene knockout or gene disruption, single gene expression from a plasmid) |
| 4 | MES capable AND advanced genetic tools (multiple gene insertion/deletion, substantial rewiring of metabolic flux) |
| 5 | MES capable AND advanced genetic tools AND industrial scaling (deployed in a commercial bioprocess) |

### Supplementary notes

**Supplementary note 1. Number density and turnover number of $CO_2$-fixing enzymes in autotrophic organisms.** Key enzymes in carbon fixation pathways are well characterized[36,37]. We used BRENDA (http://brenda-enzymes.org) to identify turnover number ranges for RuBisCo (Calvin cycle), ATP citrate lyase (rTCA cycle), 4-hydroxybutyryl-CoA dehydratase (Fuchs-Holo cycle), and carbon monoxide dehydrogenase (Wood-Ljungdahl pathway). We selected representative turnover numbers from Mueller-Cajar *et al.*[38] and Tcherkez *et al.*[39] for RuBisCo, Wahlund *et al.*[40] and Kim *et al.*[41] for ATP citrate lyase, Hawkins *et al.* for 4-hydroxybutyryl-CoA dehydratase[42], and Roberts *et al.*[43] for carbon monoxide dehydrogenase.

To estimate the enzyme count in cells, we used the approximate total enzyme amount in *E. coli*[44] and estimated the fraction of total cell protein comprised by key enzymes in carbon fixation pathways. We used a value of 3% for RuBisCo, ATP citrate lyase, and 4-hydroxybutyryl-CoA dehydratase based on estimates for RuBisCo by Bar-On *et al.*[45] and 9% for carbon monoxide dehydrogenase based on an estimate by Roberts *et al.*[43].

**Supplementary note 2. Supplementary note 7: $O_2$ limited production.** To determine the $O_2$-limited production rate, we simplify our model by assuming that $O_2$ is fully saturated in the bulk electrolyte and assume that aerobic respiration functions at $O_2$ concentrations >3 nM[46]. Pyruvate production in this scenario is limited primarily by the low solubility in electrolyte solutions (~176 μM at atmospheric $O_2$ partial pressure in our system).

**Supplementary note 3: An applied voltage minimum as a function of biofilm thickness.** The fundamental tradeoff between reduced activation overpotential and increased transport losses when the biofilm thickness increases implies the existence of a minimum in the applied voltage necessary to achieve a specific pyruvate production rate. For microbes fixing carbon using the rTCA cycle in our model, this minimum occurs at 85, 60, and 54 μm for 5, 10, and 15 μmol/cm$^2$/hr production rates, respectively. However, these minima are likely to be highly sensitive to the properties of specific reactors and operating conditions: at 10 μmol/cm$^2$/hr, biofilms between 35 and 94 μm thick operate at voltages within 1% of the calculated applied voltage minimum, suggesting that minute adjustments of biofilm thickness (*e.g.* via dilution rate or agitation[47,48]) are unlikely to improve efficiency at a given production rate.

**Supplementary note 4. Distribution of multi-heme cytochromes in the *Chlorobia***
As an unsequenced strain of *Prothecochloris aestruarii* has been shown to directly uptake electrons from a cathode, and, as the three *Prosthechochloris spp.* included in our dataset did not encode any of the four cyctochromes in our study (Table S6), we searched for putative cytochromes in our multi-heme cytochrome phylogeny that were from the *Chlorobia* and found two clades. Group I occurred within a clade defined by the TIGRfam TIGR04315 whose only characterized representative is a tetrathionate reductase, suggesting that these proteins may not be involved in electron transfer with an electrode. Group II claded closer to ExtA and the DmsE/MtrA/PioA/MtoA clade allowing for the possibility that this clade of *Chlorobi* proteins may be involved in extracellular electron transfer.

**Supplementary note 5. Comparing intracellular diffusion and reaction rates for substrates in carbon fixation pathways.** The rate of intracellular diffusion of substrates can be compared to the reaction rate of substrates using the Thiele modulus, defined for Michaelis-Menten kinetics by[49]:

$$\phi = \frac{r_{\text{cell}}}{3}\left(\frac{v_{\max}}{K_{\text{M}} D_{\text{eff}}}\right)^{1/2} \quad (1)$$

where $r_{\text{cell}}$ is the microbial cell radius, $v_{\max}$ is the maximum reaction rate per unit volume of catalyst, $K_{\text{M}}$ is the Michaelis-Menten constant for the enzyme, and $D_{\text{eff}}$ is the effective diffusion coefficient of the substrate inside the microbial cell. Using values tabulated in the main text (Table 3), with $K_{\text{M}} = 0.34$ mM[39], we calculate $\phi = 9.8 \times 10^{-3}$ for carbon fixation with $CO_2$ as the substrate and RuBisCo as the rate-limiting enzyme.

This low value indicates that the carbon fixing reaction is limited intracellularly by enzymatic catalysis and not by intracellular diffusion; in other words, energy carriers and $CO_2$ have complete and effectively immediate access to intracellular enzymes once generated at or delivered to the cell surface. Therefore, the carbon fixing reaction can be treated as occurring at the cell surface without any loss in model validity.

We note that this assumption becomes less valid as the turnover number, intracellular enzyme concentration, or cell radius increases and therefore caution that this assumption should be re-evaluated for larger microbes, higher turnover numbers, or increased expression of rate-limiting enzymes. In cases where this assumption is no longer valid, a model that includes an effectiveness factor for enzyme utilization can be included following, for example, the method of Vos *et al.*[50] for biofilm catalysts.

**Supplementary note 6: $CO_2$ gas-liquid mass transfer.** Typical ranges for $k_L a$ are $10^{-2}$–$10^{0}$ $s^{-1}$ [51]. We picked $k_L a = 5 \times 10^{-1}$ $s^{-1}$ to limit the conditions under which the reactor is limited by the mass transfer of $CO_2$ from the gas phase to the liquid phase. We show the impact of the mass transfer coefficient on the operating voltage in Fig. S1. The mass transfer coefficient has only a slight impact on the operating voltage for a given production rate. However, below certain values, the $CO_2$ feed rate is insufficient to match the consumption rate in the biocathode layer. Thus, the mass transfer coefficient should be maintained at values >~0.1 to ensure sufficient $CO_2$ to drive high production rates.

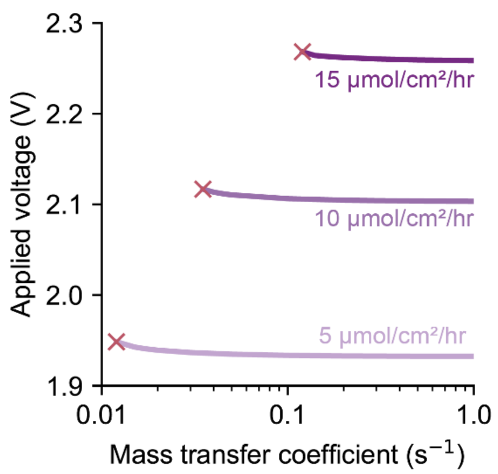

**Figure S1. Effect of $CO_2$ gas-liquid mass transfer on operating voltage.** Voltage necessary to operate the MES reactor for a 50 μm biofilm fixing carbon using the rTCA cycle as a function of the $CO_2$ gas-liquid mass transfer coefficient ($k_L a$) for different pyruvate production rates. Red crosses indicate the point at which $CO_2$ feed to the reactor is insufficient to match the consumption rate in the biocathode.

**Supplementary note 7: boundary layer thickness.** The (bio)electrochemical reduction of $CO_2$ depends strongly on the concentration polarization that develops because bicarbonate solutions are relatively weak buffers. The pH and $CO_2$ concentration at the biocathode surface and throughout the biocathode layer will vary significantly from that in the bulk electrolyte; the magnitude of this variance will depend primarily on the hydrodynamics of the electrochemical cell. Clark *et al.* measured hydrodynamic boundary layer thicknesses in an aqueous cell as function of the $CO_2$ gas flowrate and showed that increasing the flowrate decreased the boundary layer thickness but that the effect diminishes as the flowrate was increased, approaching a minimum of ~40 μm at $CO_2$ flowrates greater than ~20 sccm[52]. We assume a boundary layer of 100 μm (associated with a $CO_2$ flowrate of ~5 sccm in their system) in the main text and show how reduced boundary layer thicknesses impact operating voltages for our reactor geometry in Fig. S2. At a production rate of 15 μmol/cm$^2$hr, the total voltage increases from ~2.17 V to ~2.26 V, indicating an increased overpotential of ~100 mV. Reduced boundary layer thickness, therefore, plays an important role in MES reactor efficiency at high production rates (current densities) and is worth considering in the context of expected reactor productivity.

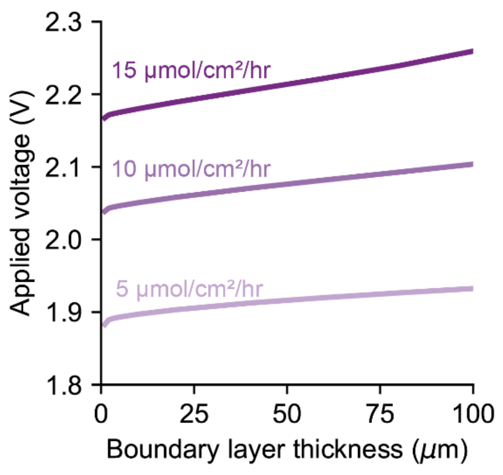

**Figure S2. Effect of boundary layer thickness on supplied voltage.** Voltage necessary to operate the MES reactor at for a 50 µm biofilm fixing carbon using the rTCA cycle as a function of the boundary layer thickness for different pyruvate production rates.

**Supplementary note 8: equilibrium potential for mtrCAB reduction.** Multiheme cytochromes typically exhibit a potential window rather than a single equilibrium potential at which electrons are reversibly exchanged with an electron (*e.g.* –400 mV to +100 mV *vs.* SHE for MtrF, an MtrC analog) corresponding to the different redox environments for each heme group within the protein[53,54]. We use –0.1 V *vs.* SHE at pH 7 (0.314 V *vs.* RHE) as a representative midpoint of the potential window for MtrCAB that remains electropositive enough to reduce the quinone pool (~–80 mV *vs.* SHE at pH 7).

**Supplementary note 9: biofilm conductivity.** The importance of biofilm conductivity in determining the productivity of biofilms in either microbial fuel cells or MES systems is the subject of ongoing debate. Some experimental studies have suggested conductivity is a critical parameter, while some modeling studies suggest the opposite[55,56]. We use a biofilm conductivity of 1 mS/cm to match that of the *Geobacter sulfurreducens* BEST strain and plot the dependency of cell voltage on biofilm conductivity in Fig. S3. Increasing conductivity above ~1 mS/cm confers only a very small reduction in total applied voltage for low production rates. However, at higher production rates (>10 µmol/cm$^2$hr), the necessary applied voltage increases rapidly for biofilm conductivities <1 mS/cm. Conductivities above 1 mS/cm have been readily achieved for *Geobacter* spp. biofilms, so synthetic strategies to increase biofilm conductivity (*e.g.* using metal or semiconducting nanowire scaffolds) are unlikely to result in significant performance enhancements in the short term, especially relative to the additional cost and complexity of fabrication.

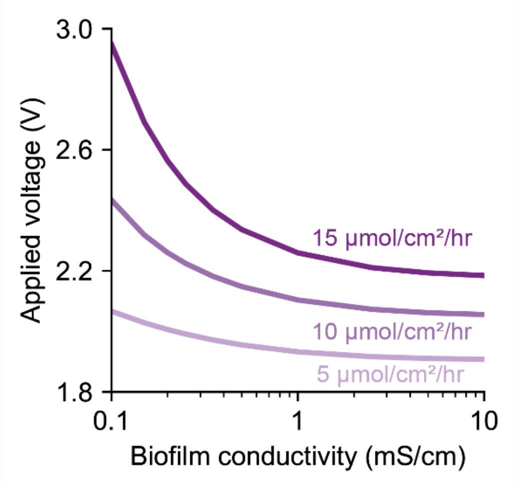

**Figure S3. Effect of biofilm conductivity on operating voltage.** Voltage necessary to operate the MES reactor 50 μm biofilm fixing carbon using the rTCA cycle as a function of the biofilm conductivity for different pyruvate production rates.

### Supplementary Figures

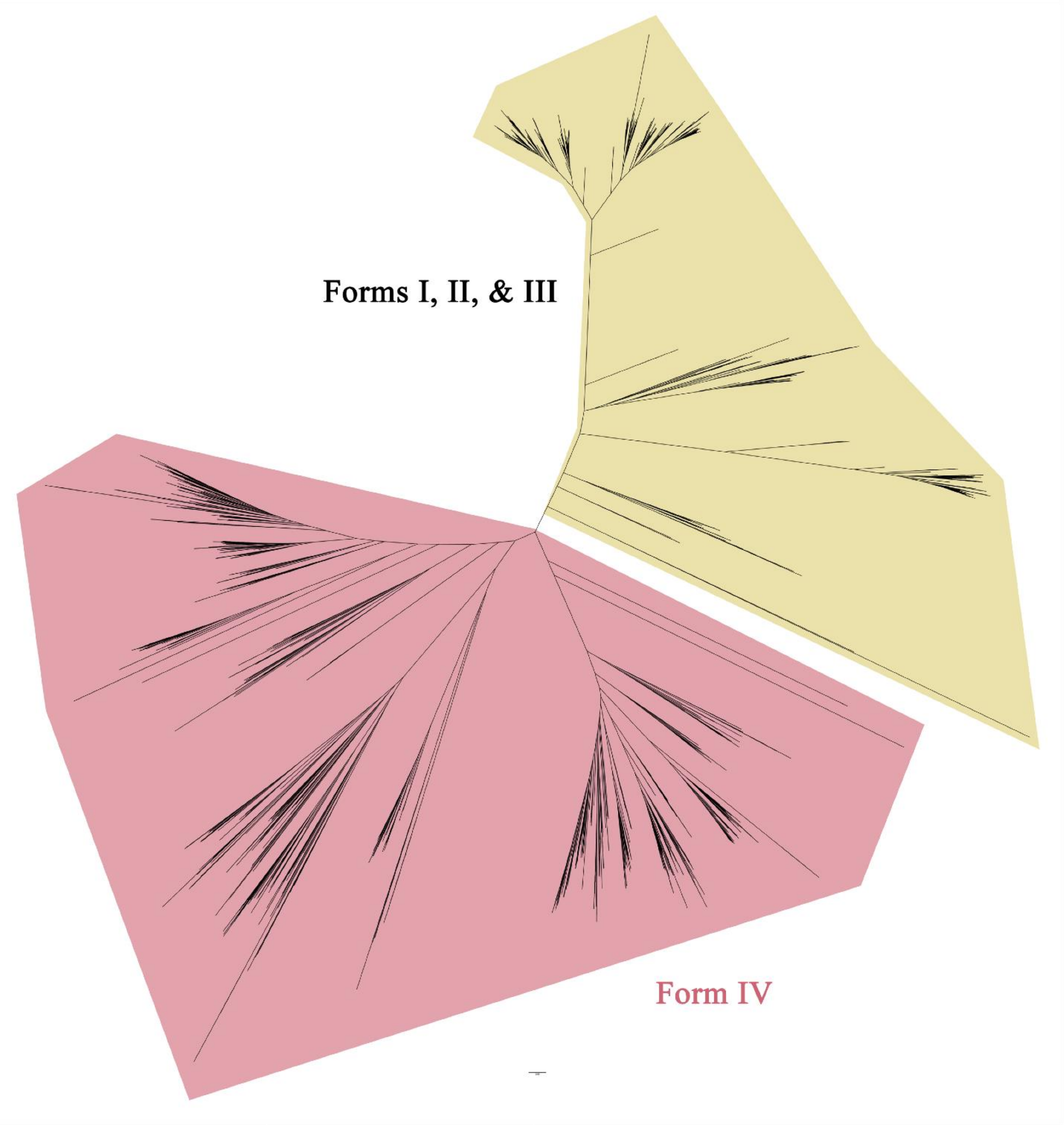

**Figure S4.** Phylogeny for the RubisCO large subunit (CcbL). Form IV (RubisCO-like; red) proteins were discarded for the analysis.

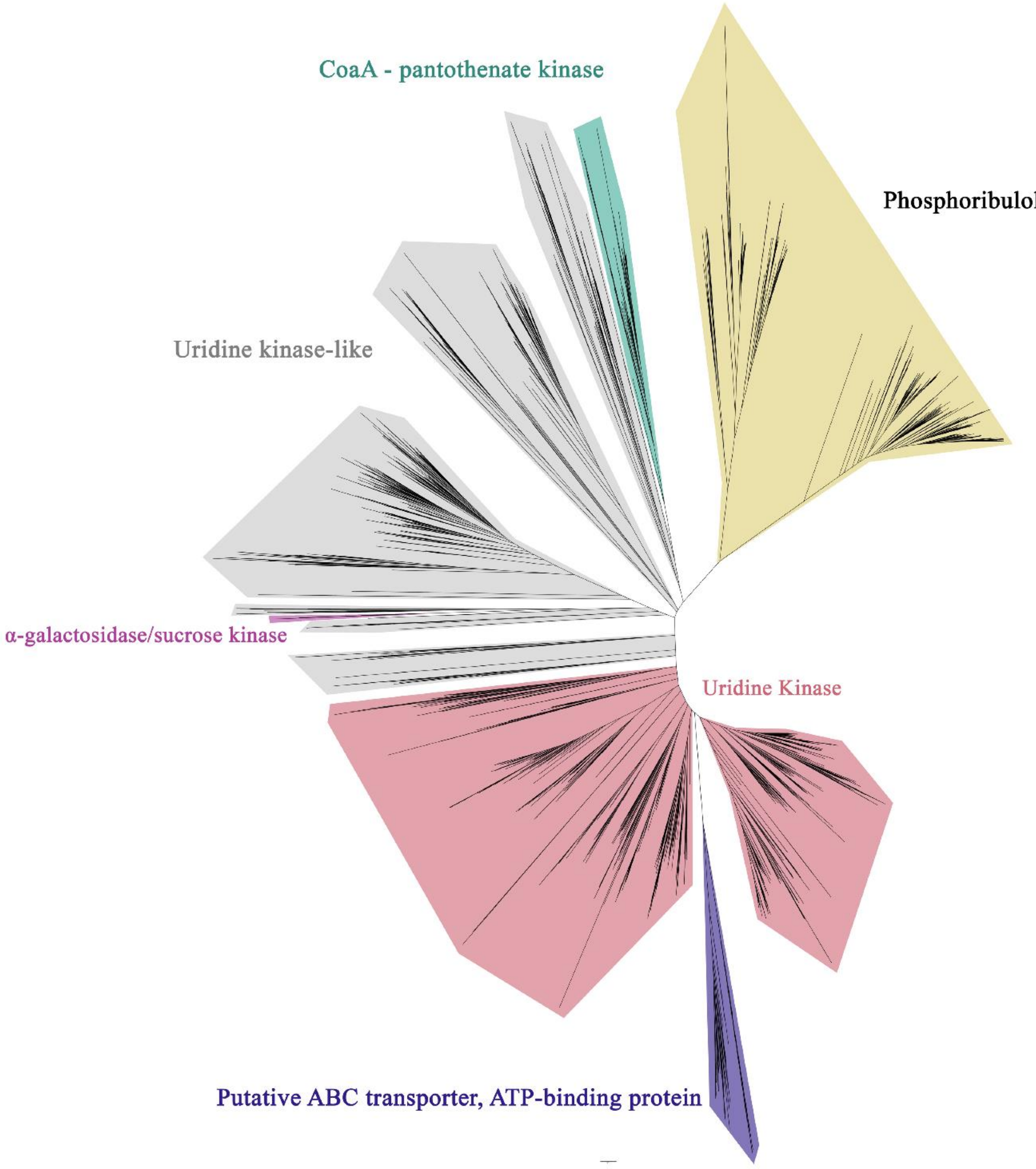

**Figure S5.** Phylogeny for phosphoribulokinase (Prk). Proteins in the Prk clade (yellow) were kept for analysis.

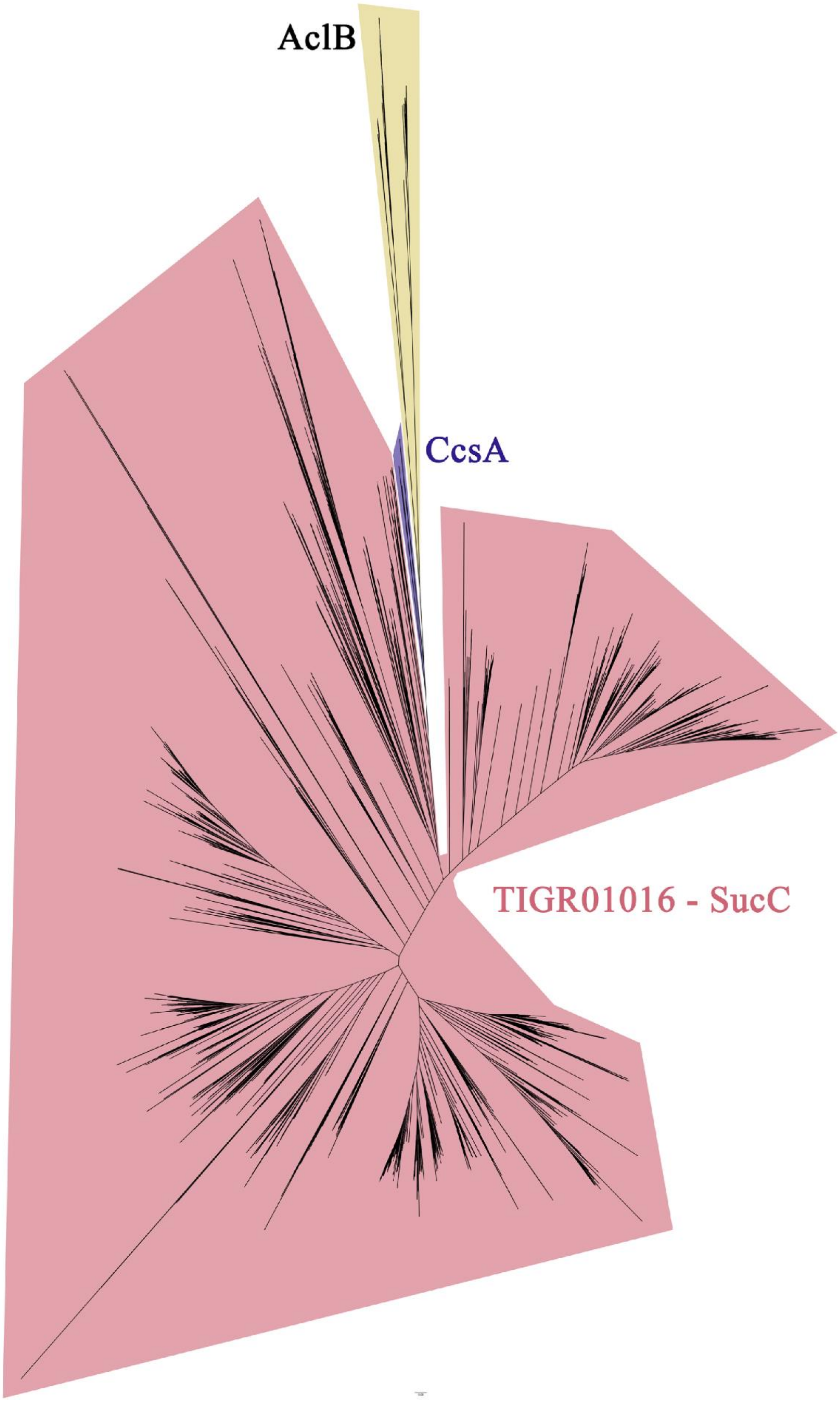

**Figure S6.** Phylogeny for AclB/CcsA. Clades annotated as AclB (yellow) and CcsA (indigo) were extracted for further analysis.

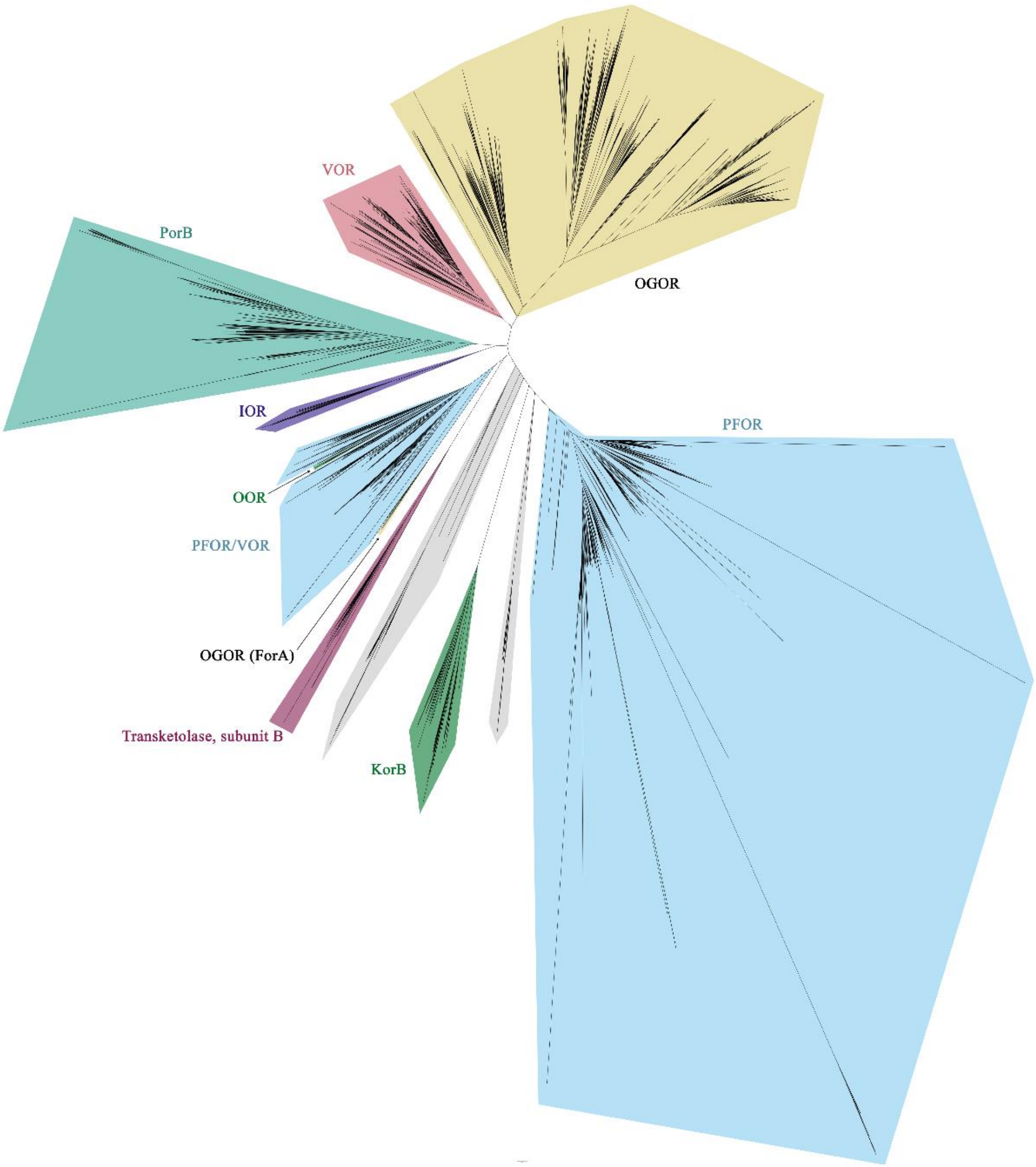

**Figure S7.** Phylogeny for 2-oxoacid oxidoreductase family proteins. Proteins within PFOR (yellow) and OGOR (blue) clades were kept for the analysis.

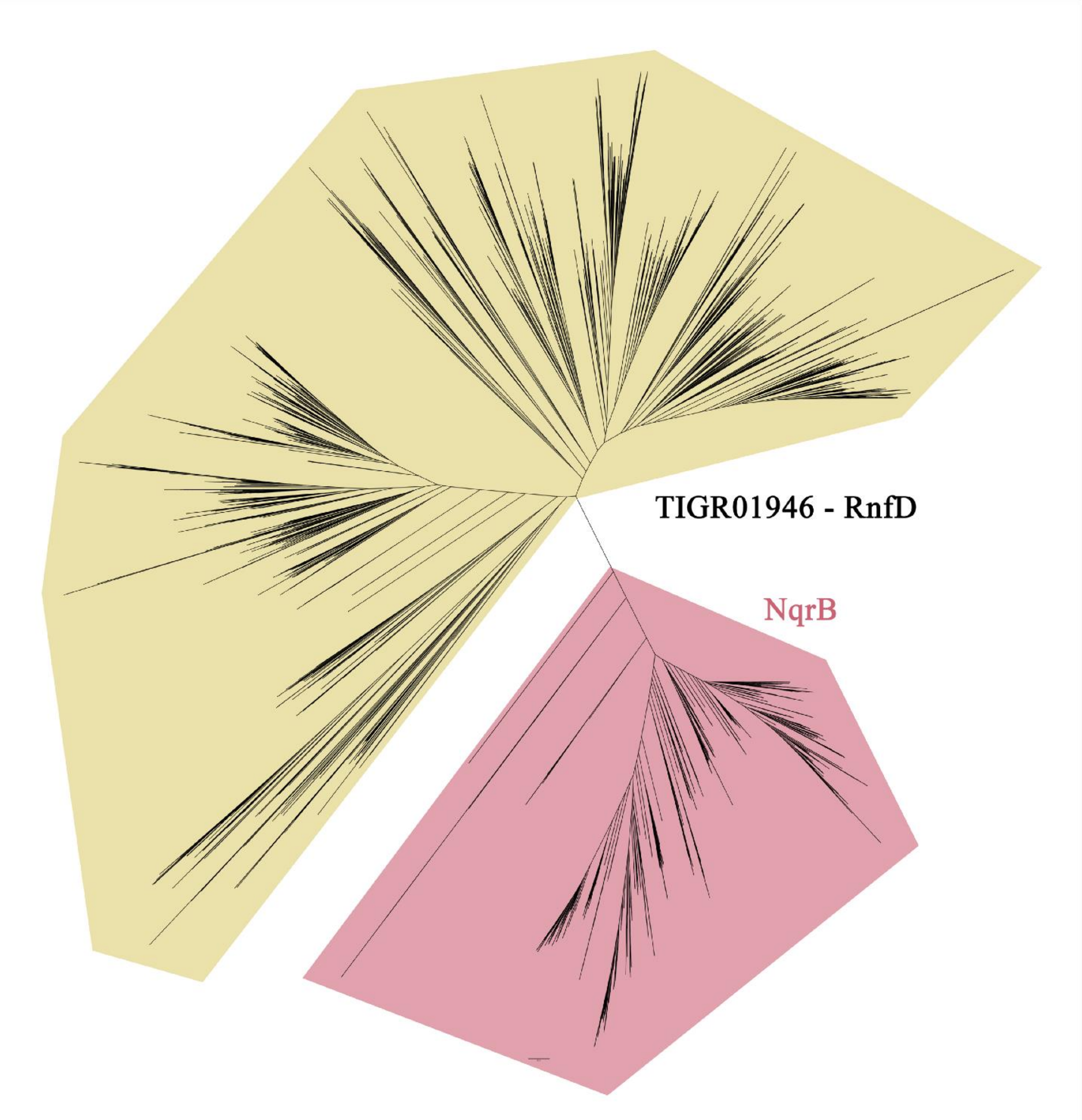

**Figure S8.** Phylogeny for RnfD. Proteins within the RnfD clade (yellow) were kept for analysis.

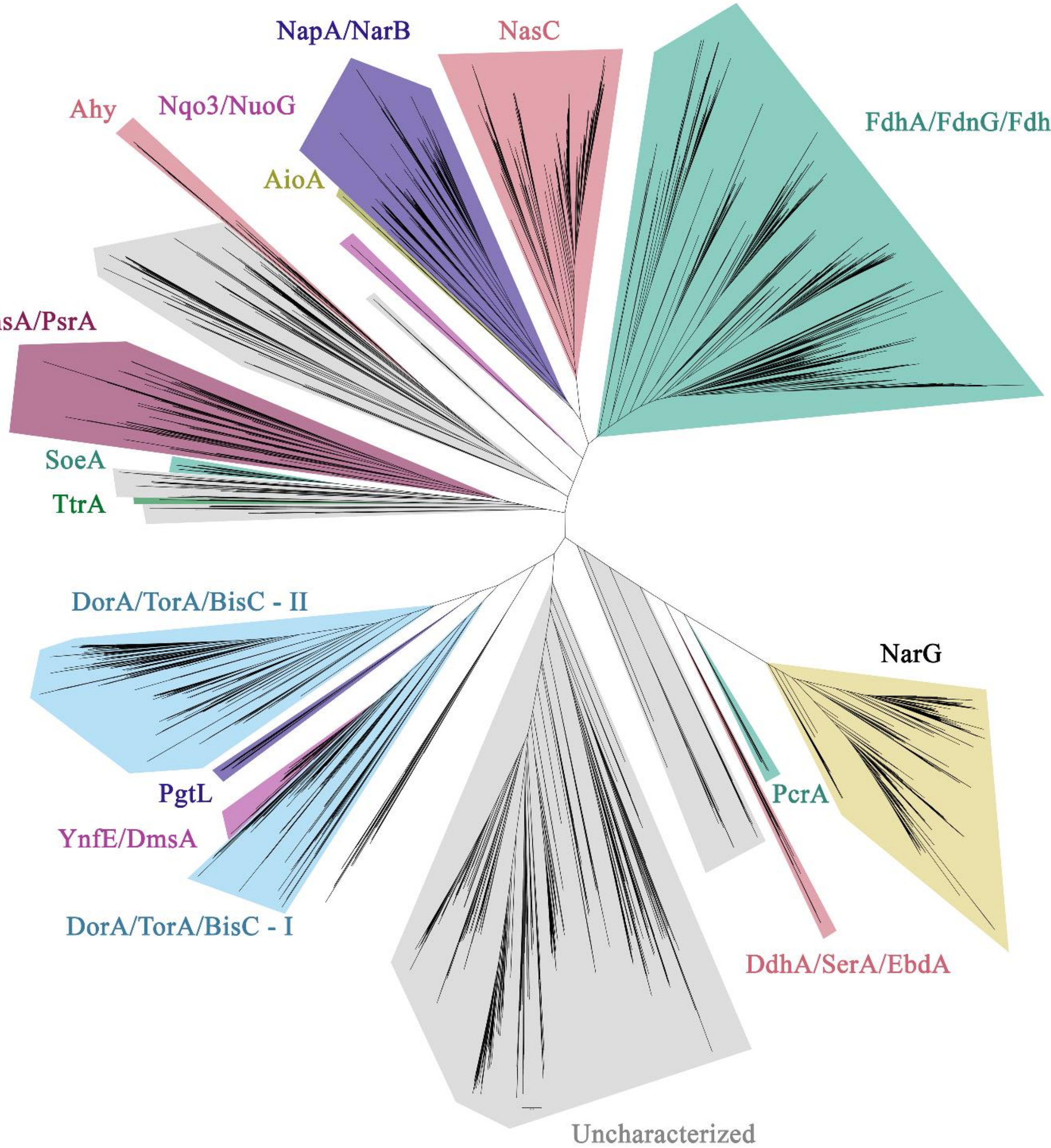

**Figure S9.** Phylogeny for molybdopterin oxidoreductase family proteins. Proteins within the NarG clade (yellow) were kept for analysis.

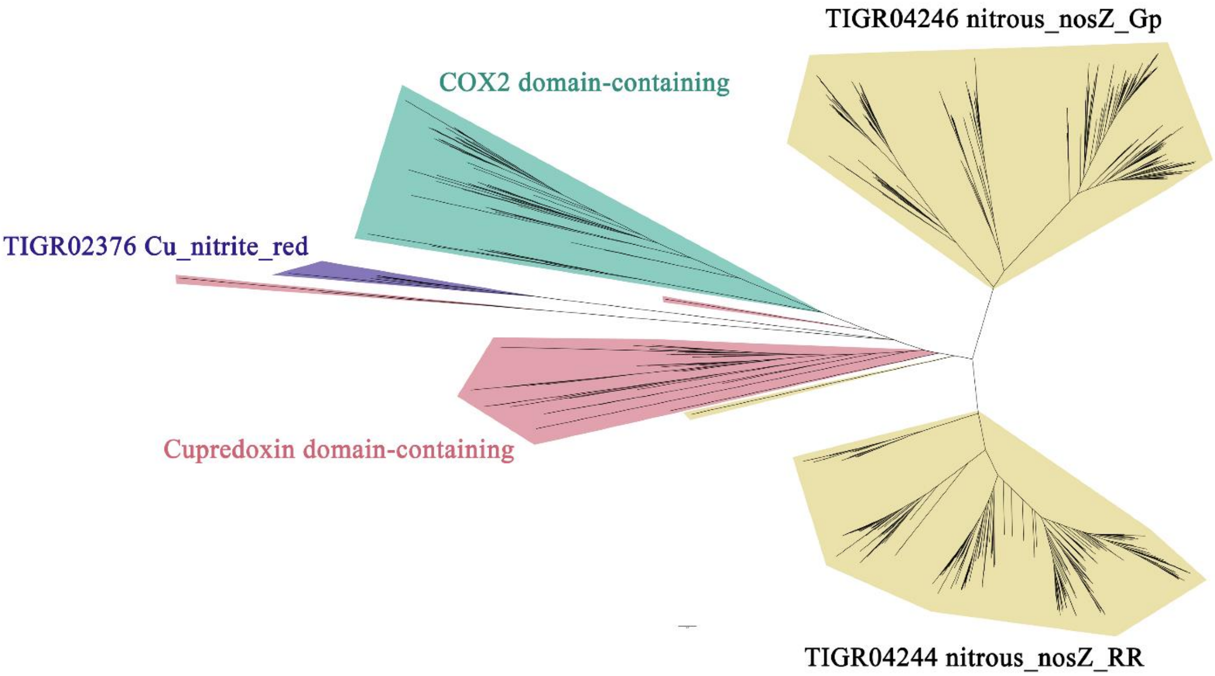

**Figure S10.** Phylogeny for nitrous oxide reductase (NosZ) and related proteins. Proteins with the NosZ clades (yellow) were kept for analysis.

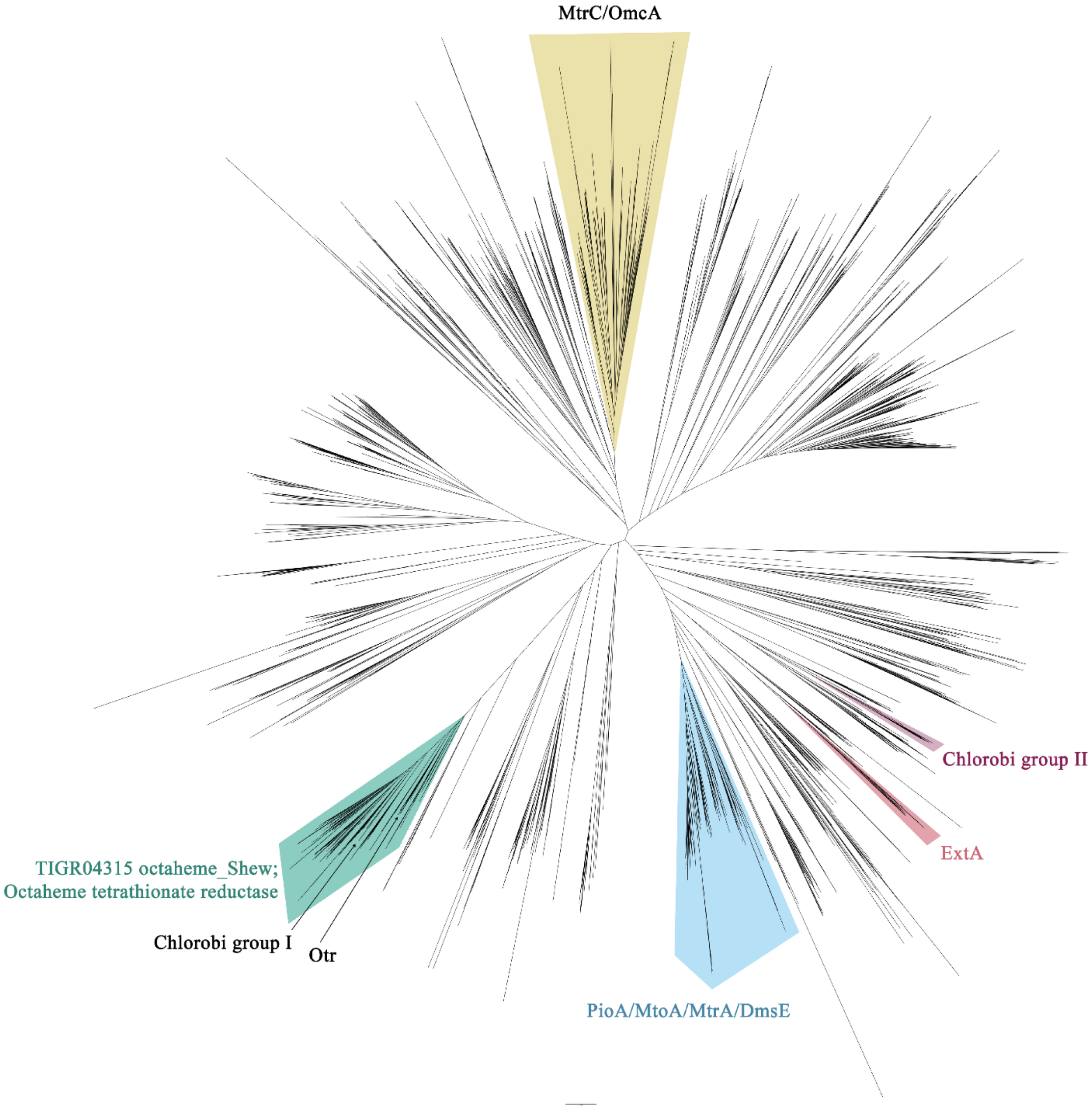

**Figure S11.** Phylogeny for multi-heme cytochromes based on an MtrC seed sequence. Sequences from the MtrC/OmcA clade (yellow), the PioA/MtoA/MtrA/DmsE clade (blue), and the ExtA clade (red) were kept for analysis. Multi-heme cytochromes from the Chlorobi were found in two clades: Chlorobi group I within the TIGRE04315 clade (teal) and Chlorobi group II (wine); these were not included in downstream analysis.

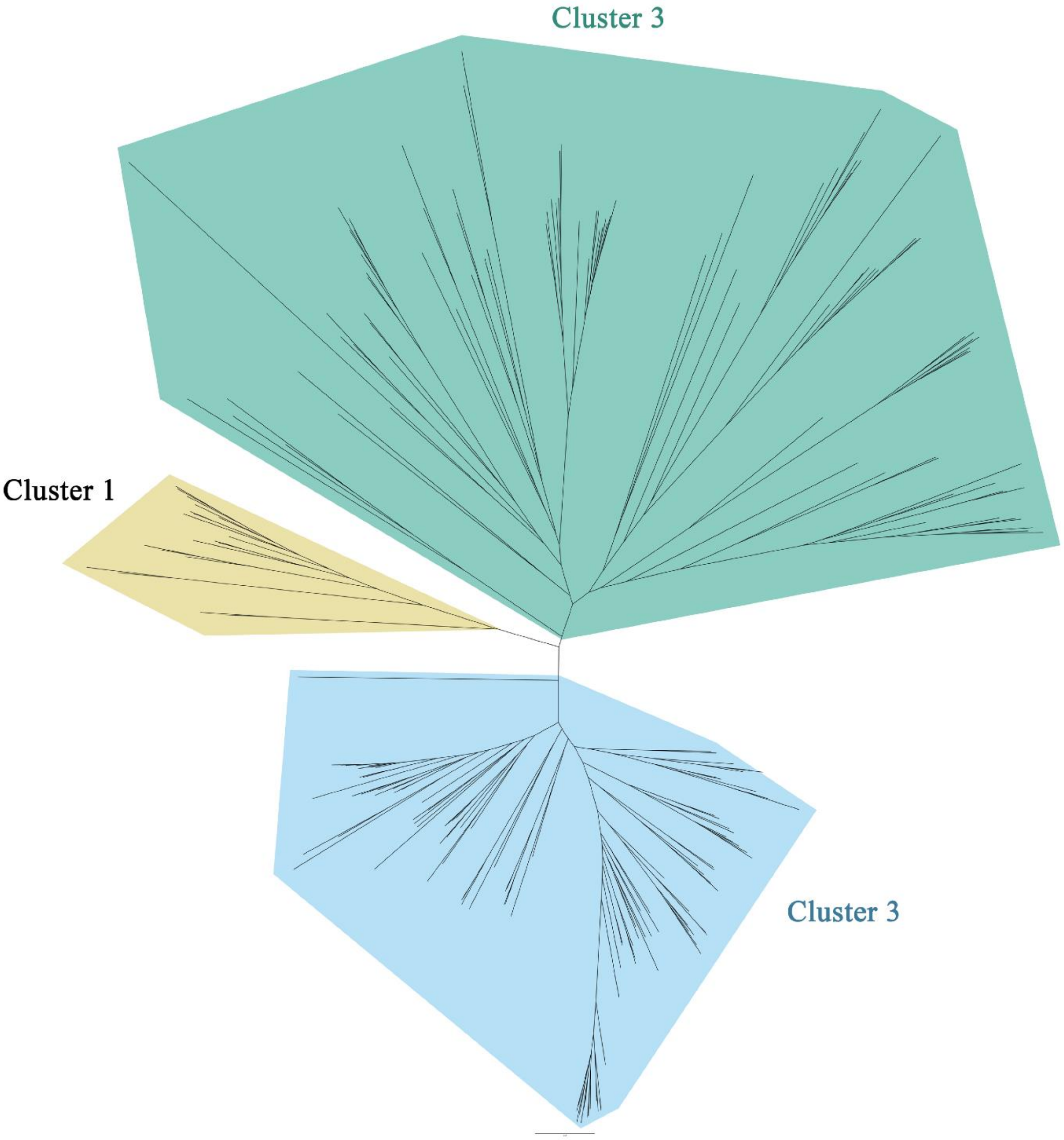

**Figure S12.** Phylogeny for Cyc2. All proteins from this tree were kept for the analysis. Topology and clustering agreed with McAllister *et al.* 2020.
